# Cytoplasmic localization of the mRNA encoding actin regulator, Serendipity-α, promotes adherens junction assembly and nuclear repositioning

**DOI:** 10.1101/2025.07.29.667534

**Authors:** Tejas Mahadevan, Lauren Renee Figard, Hasan Seede, Poonam Sehgal, Yuncong Geng, Kevin McDonald, Ido Golding, Anna Marie Sokac

**Affiliations:** Department of Cell and Developmental Biology, University of Illinois Urbana-Champaign, Urbana, IL 61801; Department of Biochemistry, Baylor College of Medicine, Houston, TX 77030; Department of Physics, University of Illinois Urbana-Champaign, Urbana, IL 61801; Center for Biophysics and Quantitative Biology, University of Illinois Urbana-Champaign, Urbana, IL 61801; Department of Microbiology, University of Illinois-Urbana Champaign, Urbana, IL 61801

## Abstract

The subcellular localization of mRNAs is conserved from prokaryotes to humans. In *Drosophila* embryos ∼70% of mRNAs localize to specific sites in the cytoplasm, but the functional significance of this mRNA localization is largely unknown. During the process of embryo cellularization, mRNA encoding Serendipity-α (Sry-α), an actin filament (F-actin) binding protein, moves apically, concentrating near centrosomes. Transport is mediated by the Egl/BicD/Dynein complex and requires two stem loops in the 3’UTR of the mRNA, which serve as localization signals. mRNA localization is dispensable for Sry-α function at cleavage furrows in early cellularization but is necessary for repositioning nuclei in late cellularization. Sry-α protein promotes assembly of cortical F-actin and apical spot adherens junctions (AJs) in late cellularization, and these AJs contribute to nuclear repositioning. We suggest that mRNA localization restricts cytoskeletal functions in late cellularization to regulate nuclear repositioning in preparation for the tissue morphogenesis events that immediately follow.

## Main

Both prokaryotes and eukaryotes exploit the subcellular localization of mRNAs to properly localize proteins and, consequently, promote cellular function ^1–3^. For example, in bacteria where transcription and translation are coupled, mRNAs encoding membrane proteins are transcribed near the inner membrane to facilitate efficient protein insertion there ^4^. Similarly, in eukaryotes, localized mRNAs can support local translation and immediate action of encoded proteins in a restricted area, as observed for β-actin mRNA which concentrates at the leading edge of vertebrate fibroblasts or in the growth cones of neurons to enhance cell motility ^5–8^. Alternatively, the asymmetric distribution of mRNAs encoding transcription factors can instruct subsequent cell fate decisions, as in budding yeast where mating-type switching is achieved by targeting the mRNA of ASH1 transcriptional repressor to daughter, but not mother, cells ^9^. With the introduction of multiplexing methodologies, it has become clear that large fractions of mRNAs localize, with some cells, such as the fruit fly embryo, showing localization of more than half of their transcripts ^10–13^. Yet, despite the widespread deployment of mRNA localization, its biological consequence remains unknown for most individual mRNAs.

Like mRNAs, organelles also show distinct subcellular distributions that influence cellular processes ^14^. The position of the nucleus, in particular, is critical to cell function. Asymmetrically positioned nuclei serve as hubs to ensure the proper organization and polarity of other sub-cellular systems, such as microtubules, ER and golgi ^15–17^. What’s more, nuclei can move to benefit activities of the cell. As the largest and most rigid organelle within cells, nuclei can be positioned to modulate cell shape and even optimize mobility ^18^. For example, in motile cells the nucleus can be positioned to “drill” or push through confined interstitial spaces; while in skeletal muscle, nuclei move to the cellular periphery to avoid mechanical interference with the contractile apparatus during muscle contraction ^19–22^. At the scale of tissues, such as highly proliferative epithelia in development, nuclei localize in a pseudostratified pattern, alternating in apical to basal position, to allow dense cell packing, and these nuclei must move apically just before mitosis to maintain epithelial integrity during cell division ^23–27^. Nuclear position in epithelia can also impact gene expression and cell fate specification because nuclei positioned in closer proximity to the tissue surface preferentially respond to extracellular cues and metabolites ^28–33^. Failures in nuclear positioning lead to numerous pathologies, underscoring the essential contribution that nuclear position makes to normal cellular function ^14^.

Nuclear position, even when it appears fixed, is actively achieved by a balance of pushing and pulling forces acting on the nucleus ^18,21^. Investigations into specific mechanisms of nuclear positioning have revealed the essential “toolbox”, which includes all the cytoskeletal polymers: actin filaments (F-actin), microtubules and intermediate filaments ^34^. F-actin and microtubules can push or pull nuclei through their own polymerization and depolymerization dynamics, by generating sweeping cytoplasmic flows, or via their motor proteins, Myosin, Kinesin and Dynein ^23,34–36^. Cytoskeletal forces and linkage can be directly conveyed to nuclei via large complexes that span the nuclear envelope and are composed of paired KASH and SUN proteins ^37^. On the outside of the nucleus KASH proteins interact with F-actin, Kinesin, Dynein or Plectin. Inside the nucleus, SUN proteins anchor to the nuclear lamina to efficiently transmit forces exerted by the cytoskeleton ^18,35,37^. Positioning may be further tuned by the physical properties of the nucleus itself, and by connectivity between the nucleus and other organelles or the cell surface ^27,36,38–40^.

While studying the mechanisms of nuclear positioning has successfully revealed these common themes, knowledge gaps remain in understanding how nuclear position and movement is regulated. Here we show that apically positioned nuclei in *Drosophila* embryos move basally during cellularization. This nuclear repositioning is dependent on the apical localization of an mRNA encoding the F-actin regulator, Serendipity-α (Sry-α). Sry-α promotes the assembly of cortical F-actin and apical spot adherens junctions (AJs), which contribute to nuclear repositioning. Thus, we reveal that the polarized localization of *sry-α* mRNA serves as an upstream regulator of nuclear repositioning in the embryo.

## Results

### *sry-α* mRNA and protein localize apically in late cellularization

The *Drosophila* embryo develops as a syncytium through its first 13 nuclear division cycles. Then, at the onset of the 14^th^ nuclear cycle, plasma membrane furrows simultaneously invaginate around each of ∼6000 cortically anchored nuclei in a process called cellularization. At the end of cellularization the resulting mononucleate cells polarize and become adherent, comprising an epithelial sheet that encases the embryo ^41^. Immediately following cellularization, the epithelial sheet initiates patterned folding as gastrulation begins ^42^.

Cellularization requires maternally contributed proteins as well as five zygotically expressed F-actin regulators ^43^. Among these regulators, Serendipity-α (Sry-α) represents a novel clade of the Vinculin/ α-Catenin Superfamily and stabilizes F-actin at invaginating furrow tips ^44^. While examining the transcriptional dynamics of *sry-α*, we noticed that its mRNA localization changes from early to late cellularization. We followed *sry-α* mRNA dynamics in live embryos using the MS2-MCP labeling system and confocal imaging ^45^. We collected Z-stacks which encompass the embryo surface and adjacent nuclei. In Z- and corresponding XZ-projections, *sry-α* mRNA (*sry-α* mRNA_MS2) localized to distinct foci inside nuclei at early cellularization (Fig. 1a, 5 minutes post-cellularization onset). The number of foci inside individual nuclei peaked 10-20 minutes after the onset of cellularization (Fig. 1c, Early). These foci are indicative of transcription occurring at the *sry-α* gene ^46^, consistent with *sry-α*’s high level of expression in early cellularization ^44^. Expression of *sry-α* is downregulated by mid-to-late cellularization ^44^, and accordingly, foci number inside the nuclei decreased (Fig. 1c, Late). As transcriptional foci disappeared with the progression of cellularization, we unexpectedly saw an increasing concentration of *sry-α* mRNA outside and apical to the nuclei (Fig. 1b, 50 minutes post-cellularization onset). By late cellularization, *sry-α* mRNA appeared in prominent apical cytoplasmic foci above the nuclei (Fig. 1b,c). These apical foci remained through the end of cellularization until gastrulation began (Fig. 1c). To validate the localization of *sry-α* mRNA over the course of cellularization, we used RNA fluorescence *in situ* hybridization (RNA-FISH) to detect endogenous *sry-α* mRNA. RNA-FISH also showed *sry-α* mRNA as foci inside nuclei at early cellularization and as apical foci above nuclei at late cellularization (Fig. 1d,e).

**Fig. 1.**
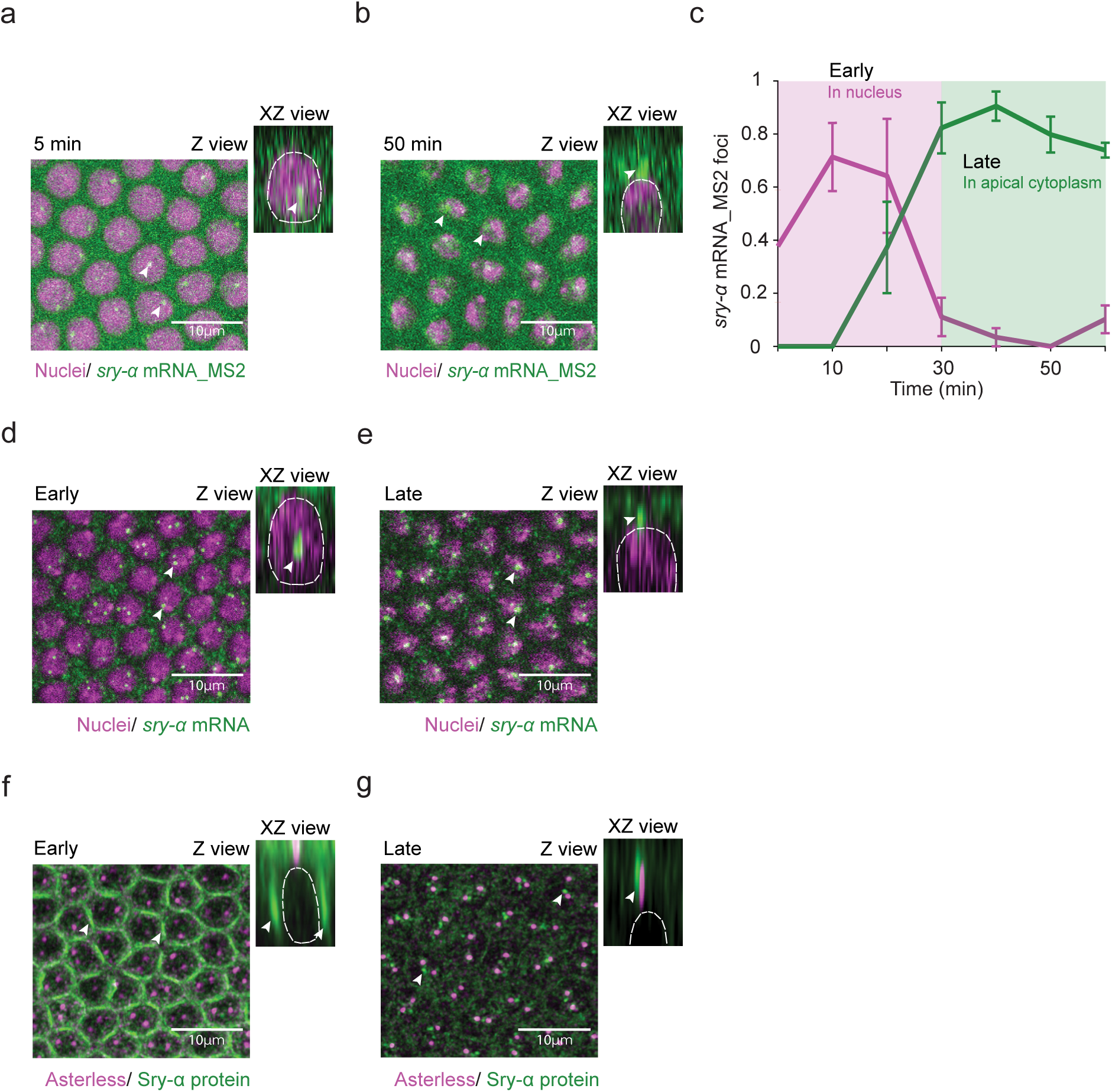
*sry-α* mRNA and protein concentrate in apical foci as cellularization proceeds. **a,b**, From time-lapse imaging, Z and XZ views show *sry-α* mRNA_MS2 (MCP-GFP; green) and nuclei (Histone-mCherry; magenta) at 5 or 50 minutes (min) post-cellularization onset. *sry-α* mRNA localizes to transcriptional foci inside nuclei at 5 minutes (**a**, arrowheads) and to apical foci in the cytoplasm above nuclei at 50 minutes (**b**, arrowheads). **c**, Number of *sry-α* mRNA_MS2 foci inside (magenta) or apical to (green) individual nuclei over the course of cellularization (each data point represents n = 4 embryos, ∼50 nuclei per embryo; mean ± s.e.m.). **d**,**e**, From RNA-FISH imaging, Z and XZ views show endogenous *sry-α* mRNA (green) and nuclei (Hoescht; magenta) at early or late cellularization. *sry-α* mRNA localizes to transcriptional foci inside nuclei at early cellularization (**d**, arrowheads) and to apical foci above nuclei at late cellularization (**e**, arrowheads). **f**,**g**, From immunofluorescence imaging, Z and XZ views show apical centrosomes (Asterless; magenta) and Sry-α protein (green) at early or late cellularization. Sry-α protein localizes to ingressing furrows at early cellularization (**f**, arrowheads) and to foci near centrosomes at late cellularization (**g**, arrowheads). **d**-**g**, Data collected from wild-type (OreR) embryos. **a**,**b**,**d**,**e**,**f**,**g**, In Z views, scale bars = 10 μm; In XZ views, white dashed line shows periphery of a nucleus. All views are projections from confocal stacks. **d**-**g**, Furrow lengths less than or greater than 5 μm indicate early or late cellularization, respectively.

To determine whether Sry-α protein also localizes apically in late cellularization, we did immunofluorescence staining. Per prior observations ^44,47^, Sry-α protein was enriched at invaginating furrow tips during early cellularization, when furrow lengths were short (0-5 µms; Fig. 1f). However, by late cellularization, Sry-α furrow enrichment was lost. Instead, we saw apical foci of Sry-α protein just above the nuclei, and coincident with the centrosomal marker, Asterless (Fig. 1g) ^48^. We suggest that *sry-α* is transcribed in the nucleus in early cellularization, and then both the mRNA and protein concentrate in a cytoplasmic position apical to nuclei during mid-to-late cellularization.

### Dynein and its adaptor BicD/Egl are involved in *sry-α* mRNA localization

The apical foci of *sry-α* mRNA are positioned close to the centrosomes that also sit just apical to nuclei in cellularizing embryos ^49^. This suggests that *sry-α* mRNA may use Dynein-mediated trafficking to move apically towards microtubule minus ends located near the centrosomes. To test this, we performed RNA-FISH in embryos following treatment with Dynein inhibitor Ciliobrevin D ^41,50^. In control embryos, treated with DMSO carrier alone (Control), apical foci of *sry-α* mRNA were obvious in late cellularization (Fig. 2a,c). However, in embryos treated with Ciliobrevin D, *sry-α* mRNA apical localization was disrupted, with few, if any apical foci apparent by late cellularization (Fig. 2b,c). Thus, the apically-directed transport and/or anchoring of *sry-α* mRNA requires Dynein activity.

**Fig. 2.**
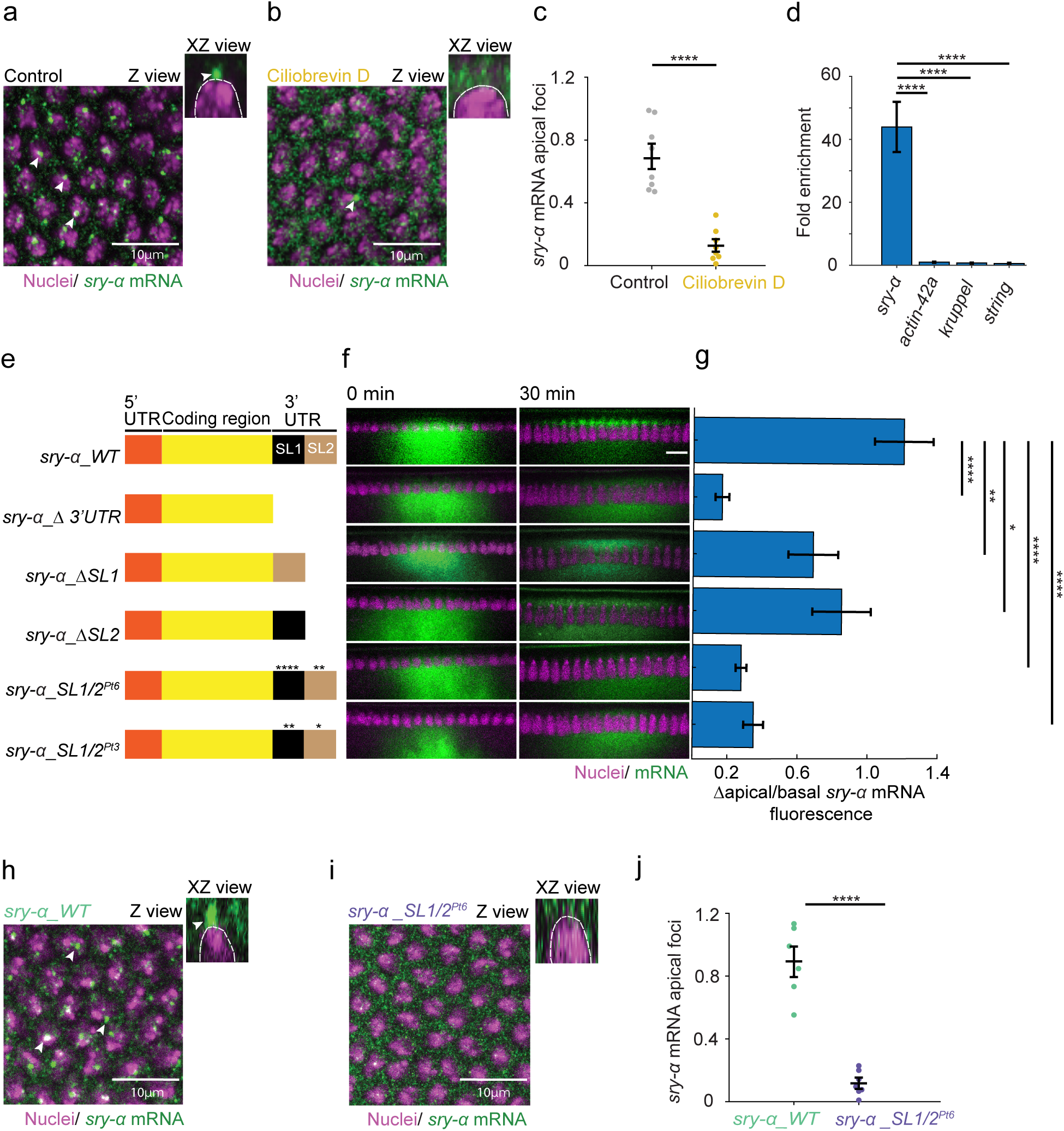
Stem loops in the *sry-α* 3’ UTR are required for Dynein/BicD/Egl mediated localization of the mRNA to apical foci. **a**,**b**, From RNA-FISH imaging, Z and XZ views show endogenous *sry-α* mRNA (green) and nuclei (Hoescht; magenta) in DMSO (**a**, Control) and Ciliobrevin D (**b**) treated embryos. *sry-α* mRNA localizes to apical foci above most nuclei in Control embryos (**a**, arrowheads), but fails to localize to the same extent after Ciliobrevin D treatment (**b**, arrowhead). **c**, Number of *sry-α* mRNA foci apical to individual nuclei at late cellularization (each data point represents average number of foci per nucleus in 1 embryo, ∼87 nuclei per embryo; n = 8 and n = 6 embryos for Control and Ciliobrevin D treatment, respectively). **d**, Fold enrichment of *sry-α* PCR product versus negative controls (*actin 42A*, *kruppel*, *string*) from BicD immunoprecipitates (n = 3 biological replicates). **e**, Schematic shows construct design for *in vitro* transcribed *sry-α* mRNAs (WT, wild-type; SL, stem loop; Pt, point mutations). Asterisks indicate point mutations. **f**, From time-lapse imaging, cross-sections show injected *sry-α* mRNA (green) and nuclei (Histone-mCherry, magenta). Identity of the *sry-α* mRNA is indicated by aligned construct in (**e**). Imaging started immediately after injection and proceeded for 30 minutes (0 min and 30 min, respectively). **g**, Change in apical to basal fluorescence from 0 to 30 minutes for the *sry-α* mRNA indicated by aligned construct in (**e**) (n ≧ 5 embryos per construct). **h**, **i**, From RNA-FISH imaging, Z and XZ views show *sry-α* mRNA (green) and nuclei (Hoescht; magenta). *sry-α* mRNA from a wild-type transgene (**g**, *sry-α_WT*) localizes to apical foci (arrowheads), while *sry-α* mRNA from a transgene with six point mutations in the 3’UTR stem loops (**h**, *sry-α_SL1/2^Pt6^*) fails to localize. **i**, Number of *sry-α* mRNA foci apical to individual nuclei at late cellularization (each data point represents average number of foci per nucleus in 1 embryo, ∼87 nuclei per embryo; n = 6 and n = 7 embryos for *sry-α_WT* and *sry-α_SL1/2^Pt6^*, respectively). **a**,**b**,**g**,**h**, Images collected from embryos at late cellularization, with furrow lengths > 5 μ m. In Z views, scale bars = 10 μm; In XZ views, white dashed line shows periphery of a nucleus. All views are projections from confocal stacks. **f**, Scale bar = 10 μm. All images are single plane confocal cross-sections. **c**,**j**, Horizontal lines indicate mean ± s.e.m.. **d**,**g**, Bars indicate mean ± s.e.m.. **c**,**d**,**j**, ****p<0.00005, Student’s t test **g,** *p<0.05, **p<0.005, ****p<0.00005, one-way ANOVA

One mechanism of Dynein-mediated mRNA transport in *Drosophila* utilizes the Dynein adaptor complex composed of proteins Bicaudal-D (BicD) and Egalitarian (Egl) ^51,52^. Previously, in RNA immunoprecipitation assays prepared from overnight embryo collections it was shown that Egl interacts with *sry-α* mRNA ^52^. To confirm an interaction between the BicD/Egl complex and *sry-α* mRNA, we did BicD immunoprecipitation from hand-staged embryos at mid-to-late cellularization, and measured levels of associated mRNA by QPCR. We found that *sry-α* mRNA is highly represented among the BicD interacting mRNAs, showing 40-fold enrichment over negative controls that do not localize apically, nor interact with Egl (*actin-42a*, *kruppel* and *string*; Fig. 2d) ^53^. We have shown previously that BicD and Dynein heavy chain concentrate near centrosomes in cellularizing embryos ^54^. Thus, we suggest that *sry-α* mRNA associates with the BicD/Egl complex, which tethers the mRNA to Dynein for apical localization.

### Stem loops in the 3’UTR serve as localization signals for *sry-α* mRNA

The localization of mRNAs is commonly mediated by cis-acting sequences that interact with RNA binding proteins and direct specific subcellular distributions ^2,3^. These localization sequences can occur anywhere in a transcript and often generate stem loops and secondary RNA structures ^51,55–58^. To investigate whether cis-acting sequences are involved in the apical localization of *sry-α* mRNA, we generated reporter constructs informed by RNA folding algorithms that predicted two large stem loops in the 3’UTR of the *sry-α* transcript (Fig. 2e). Constructs were used as template to synthesize fluorophore-labeled mRNAs *in vitro.* These labeled mRNAs were injected into live cellularizing embryos in a region basal to the nuclei and imaged over time to assess apical mobility (Fig. 2f). Per expectations for this assay ^53^, a positive control, *bicoid* mRNA (*bicoid_WT*), localized apically; while a negative control, *string* mRNA (*string_WT*), exhibited no specific localization and diffused over time (Extended Data Fig. 1). For full-length *sry-α* mRNA (*sry-α_WT)* apical movement started within 2 minutes after imaging started, and by 30 minutes, the majority of signal was concentrated apical to nuclei (Fig. 2e-g). This result confirmed that the *in vitro* synthesis and injection approach recapitulates endogenous *sry-α* mRNA localization.

Next, we generated a hybrid mRNA in which the 3’UTR of the negative control *string* was replaced by the 3’UTR of *sry-α* (*string_sry-α*). Unlike *string_WT*, the *string_sry-α* mRNA moved apically (Extended Data Fig. 1), showing that the *sry-α* 3’UTR is sufficient to drive apical localization. To ask if the 3’UTR is necessary for apical localization of *sry-α* mRNA, we generated a full 3’UTR deletion (*sry-α_Δ 3’UTR*). This mRNA completely failed to localize (Fig. 2e-g). To further map the signal sequences in the *sry-α* mRNA, we generated partial 3’ UTR deletions that eliminated either the first or second stem loop (*sry-α_ΔSL1* or *sry-α_ΔSL2*, respectively). Both *sry-α_ΔSL1* and *sry-α_ΔSL2* showed partial apical localization, resulting in approximately half the enrichment of the wild-type construct (Fig. 2e-g), suggesting that the two stem-loops contribute additively to apical *sry-α* mRNA localization. As a final test of the importance of the stem loops, we generated mRNAs with either six (*sry-α_SL1/2^Pt^*^6^) or three (*sry-α_SL1/2^Pt3^)* point mutations designed to disrupt the base-pairing in each stem structure. Neither of these point mutant mRNAs were able to localize apically, and instead diffused away from the injection site (Fig. 2e-g). So far, our results argue that the two stem-loops in the 3’UTR are needed to direct apical localization of *sry-α* mRNA.

As a final test of the role of the stem-loops in the *sry-α* 3’UTR, we made flies that express large BAC-derived transgenes encoding either *sry-α* mRNA that is wild-type or contains the same six point mutations as in the *sry-α_SL1/2^Pt6^ in vitro* transcribed construct (*sry-α_WT* or *sry-α_SL1/2^Pt6^* flies, respectively). We verified that both *sry-α_WT* and *sry-α_SL1/*2*^Pt6^* mRNAs are expressed at equal levels in embryos during cellularization (Extended Data Fig. 2). To assay mRNA localization, we used RNA-FISH for *sry-α* in embryos expressing either transgene in a *sry-α* null background (*sry-α -/-*; Fig. 2h,i). We saw that the *sry-α_WT* mRNA localized to apical foci in late cellularization, while *sry-α_SL1/*2*^Pt6^* mRNA failed to localize (Fig. 2h-j). We conclude that the two stem-loops in the 3’UTR serve as the localization sequences to direct the apical concentration of *sry-α* mRNA.

### Apical localization of *sry-α* mRNA is not required for furrow ingression

We wanted to know if the apical localization of *sry-α* mRNA in late cellularization had any functional significance. We first considered the known role played by Sry-α protein at furrows ^43^. Several prior studies have placed Sry-α protein at furrow tips in early cellularization and reported its activity in maintaining cortical F-actin levels in that position ^44,47,59,60^. Perturbation of this F-actin in *sry-α* null mutants results in destabilization and regression of cleavage furrows between adjacent nuclei, and a consequent multinucleation phenotype ^44,47,59,60^. While Sry-α’s early protein localization and function at furrows did not obviously align with the late *sry-α* mRNA localization to apical foci, we nonetheless asked: Does apical *sry-α* mRNA localization impact furrow stability? We assayed the capacity of our *sry-α_WT* and *sry-α_SL1/*2*^Pt6^* transgenes to rescue the *sry-α* null multinucleation phenotype. These experiments were performed in fixed embryos so that RNA-FISH could be used for genotyping combined with immunofluorescence for furrow detection (FISH-IF). In wild-type embryos (OreR) all furrows, as detected by Peanut/Septin staining, were intact and no multinucleation was observed (Fig. 3a). In *sry-α -/-* embryos, the multinucleation phenotype was fully penetrant, with 100% of embryos showing mild to severe regression of furrows and multinucleation (Fig. 3b). In *sry-α -/-* embryos expressing the *sry-α_WT* transgene, the multinucleation phenotype was fully rescued in all embryos (Fig. 3c). Similarly, *sry-α -/-* embryos expressing the *sry-α_SL1/*2*^Pt6^* transgene were completely rescued (Fig. 3d), even though these embryos lack apically localizing *sry-α* mRNA (Fig. 2i,j). Therefore, apical localization of *sry-α* mRNA is not necessary for the furrow function of Sry-α in early cellularization.

**Fig. 3.**
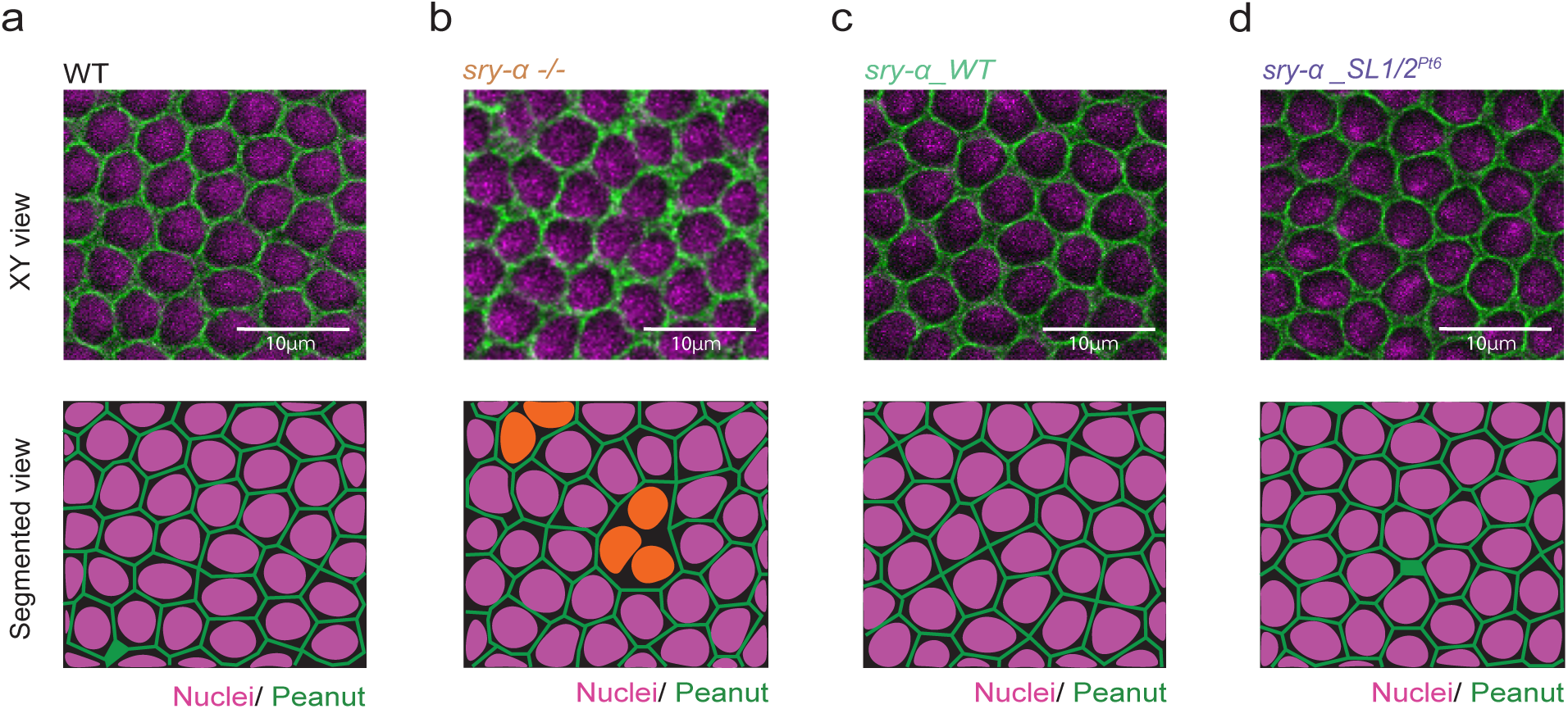
A mislocalizing *sry-α* mRNA rescues the furrow regression phenotype in *sry-α* null mutants. **a**,**b**,**c**,**d**, From FISH-IF imaging, single plane confocal XY views show furrows (Peanut, green) ingressing between nuclei (Hoescht; magenta) in wild-type (WT, OreR, **a**), *sry-α -/-* (**b**), *sry-α -/-* plus *sry-α_WT* (**c**), and *sry-α -/-* plus *sry-α_SL1/2^Pt6^* (**d**) embryos (n ≧ 10 embryos per genotype). Bottom row shows segmented views of the corresponding XY views with nuclei highlighted (orange, **b**) in multinucleated cells where furrows regressed. Images collected from embryos at late cellularization, with furrow lengths > 5 μm. Scale bars = 10 μm.

### Apical localization of *sry-α* mRNA is required for nuclear repositioning

We reasoned that Sry-α may have an unknown role in late cellularization that does depend on its apical mRNA localization. One event that happens by late cellularization is the basally directed repositioning of cortically anchored nuclei (Fig. 4a) ^61,62^. This repositioning is critical in moving nuclei out of the way of mechanical forces that are exerted at the embryo surface during gastrulation ^63^. We imaged Histone-GFP embryos to visualize the major basal displacement of nuclei away from the embryo surface over the course of cellularization (Fig. 4b). The vitelline membrane, which is one of the shells encasing the embryo, is autofluorescent and served as a marker for the embryo surface. To test for a role for Sry-α in nuclear repositioning, we used an RNAi knockdown approach. Histone-GFP embryos were injected with double-stranded RNA (dsRNA) that strongly reduces *sry-α* transcript levels ^44^. These *sry-α* RNAi embryos showed disrupted nuclear repositioning in late cellularization compared to buffer injected controls (Fig. 4b,c). To quantify the disruption, we measured the distance between the apical tops of nuclei and the vitelline membrane (Fig. 4d, see Extended Data Figs. 3 and 4 for raw and normalized individual embryo data, Extended Data Video 1 and 2). The *sry-α* RNAi embryos consistently showed shorter nuclear distances with largest divergence from control embryos by the end of cellularization (Fig. 4d,e). Tracking individual nuclei in controls showed a steady basal migration (Fig. 4f). However, some *sry-α* RNAi injected embryos showed nuclei “popping up” and even redirecting their motion apically after having started on a basal trajectory (Fig. 4g). Some nuclei in *sry-α* RNAi embryos also showed tilting such that their long axis no longer remained perpendicular to the embryo surface as normally seen in controls (Fig. 4f,g). These results suggest that Sry-α has more than one function during cellurization: Early, Sry-α stabilizes furrows and ensures their ingression; Late, Sry-α promotes the basal repositioning of nuclei (Fig, 4) ^44,59,60^.

**Fig. 4.**
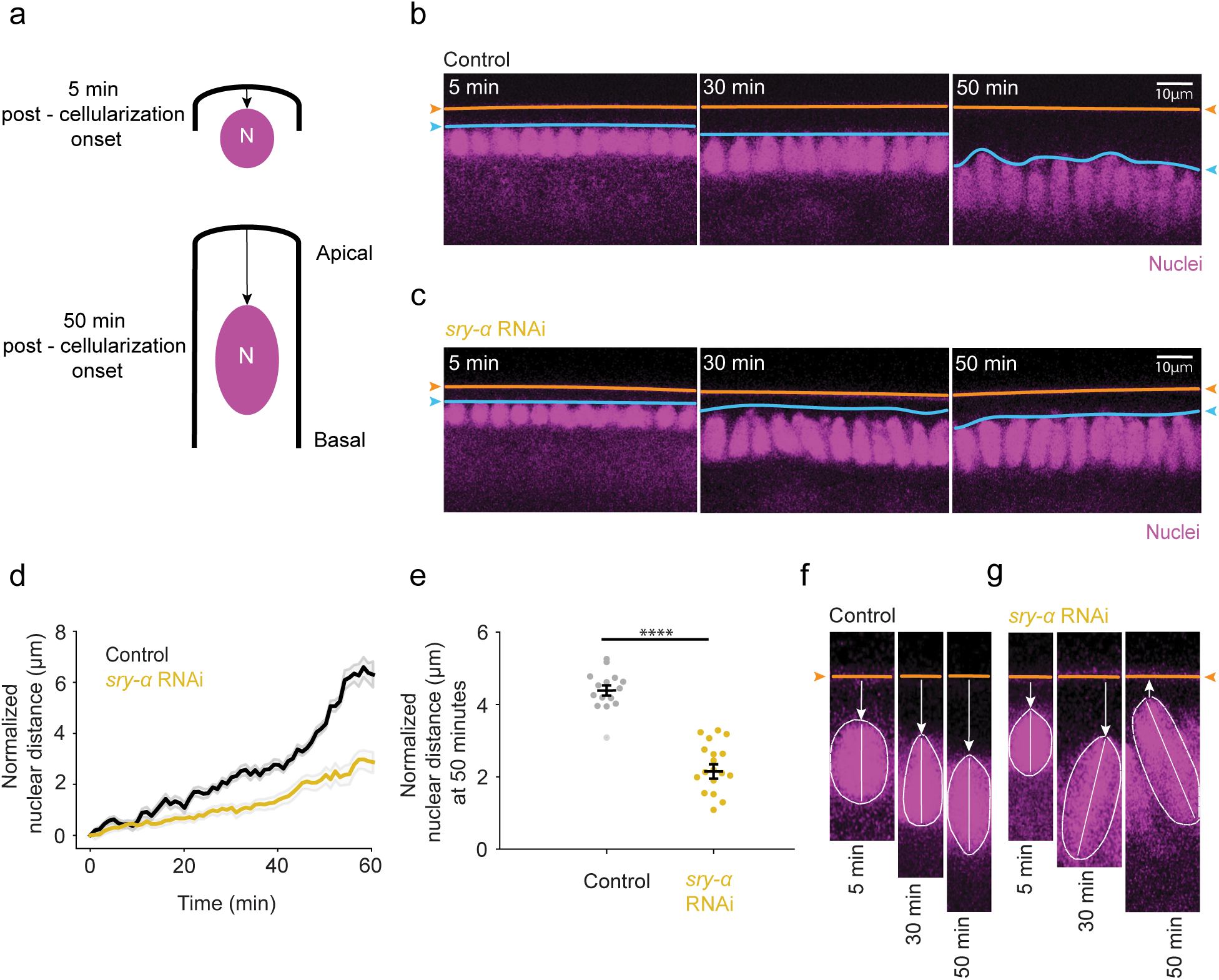
Sry-α promotes basally-directed nuclear repositioning at late cellularization. **a**, Schematic shows basal movement of nuclei from early to late cellularization (5 and 50 minutes (min) post-cellularization onset, respectively). **b**,**c**, From time-lapse imaging, cross-sections show nuclei (Histone-GFP, magenta) in buffer (Control, **b**) or *sry-α* dsRNA (*sry-α* RNAi, **c**) injected embryos at 5, 30 and 50 minutes post-cellularization onset. Orange lines and arrowheads indicate position of the vitelline membrane; blue lines and arrowheads indicate position of the apical tops of nuclei. Scale bars = 10 μm**. d**, Normalized nuclear distances for Control (black) and *sry-α* RNAi (gold) embryos over the course of cellularization (n = 5 embryos per treatment, ∼15 nuclei followed per embryo; mean ± s.e.m. demarcated in gray). **e**, Normalized nuclear distance at 50 mins post-cellularization onset in Control (black) and *sry-α* RNAi (gold) embryos (each data point represents average nuclear distance for 15 individual nuclei in n = 5 embryos per treatment). Horizontal lines indicate mean ± s.e.m.; ****p<0.00005, Student’s t test. **f**, **g**, From time-lapse imaging, higher mag cross-sections show an individual nucleus (Histone-GFP, magenta) in buffer (Control, **f**) or *sry-α* dsRNA (*sry-α* RNAi, **g**) injected embryos at 5, 30 and 50 minutes post-cellularization onset. Orange lines and arrowheads indicate position of the vitelline membrane; white arrows indicate direction of movement of the nucleus; white dashed lines show periphery of the nucleus; white solid lines indicate long axis of the nucleus. **b**,**c**,**f**,**g**, All images are single plane confocal cross-sections.

To ask if the apical localization of *sry-α* mRNA is required to support Sry-α’s role in nuclear repositioning, we tested whether our *sry-α_WT* and *sry-α_SL1/*2*^Pt6^* transgenes could rescue nuclear repositioning in a *sry-α* null background. We used FISH-IF in fixed embryos so that we could accurately genotype by FISH and determine cellularization progress based on increasing furrow length by immunofluorescence ^64^. Nuclei were detected by Hoescht staining. We first characterized the basal repositioning of nuclei in fixed wild-type (OreR) and *sry-α -/-* embryos. Fixed imaging was able to capture nuclei moving towards more basal positions over the course of cellularization in wild-type embryos (Fig. 5a). Here, “nuclear distance” was measured between the apical tops of nuclei and the plasma membrane because the vitelline membrane had to be removed to allow immunostaining. Plotting nuclear distance against furrow length showed basally-directed movement of nuclei over the course of cellularization (Fig. 5b). However, basal repositioning was impaired in *sry-α -/-* embryos, consistent with what we saw in RNAi experiments (Fig. 5d). Most nuclear distances for *sry-α -/-* embryos fell below the line of best fit determined for wild-type embryos (Fig. 5e). To further assess the extent of difference between nuclear positioning in wild-type and *sry-α -/-* embryos, we performed residual analysis for each data set (Fig. 5c,f). This analysis shows how frequently deviations of a given distance occur with respect to the best-fit line for wild-type. As expected, wild-type distances followed a normal distribution with peak probability at 0 microns (Fig. 5c). Conversely, the peak for *sry-α -/-* distances was shifted below zero, showing that nuclear distances trend shorter for *sry-α* null embryos (Fig. 5f). These results reaffirm a role for Sry-α in the basal repositioning of nuclei.

**Fig. 5.**
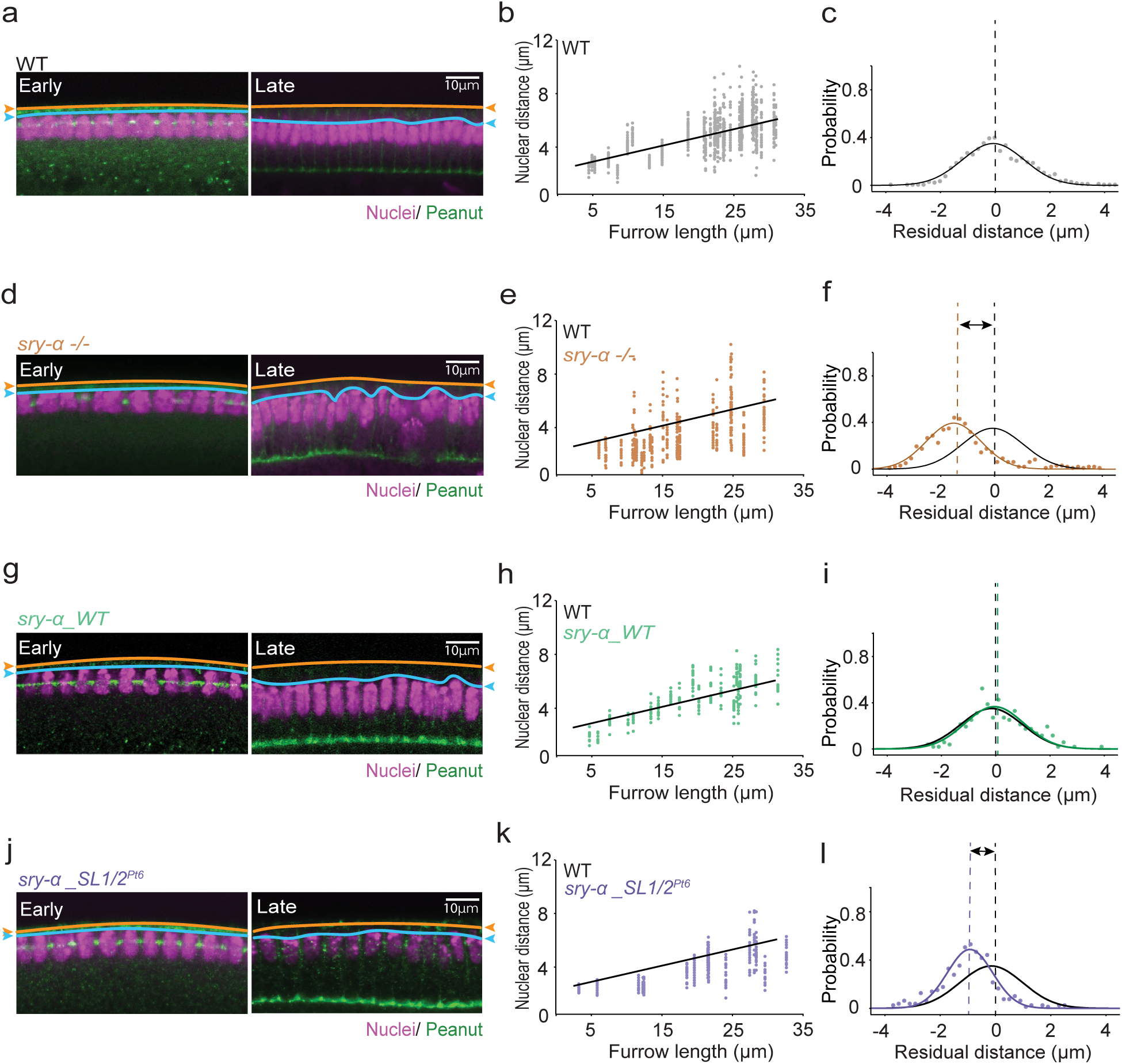
Apical localization of *sry-α* mRNA promotes nuclear repositioning in late cellularization. **a**,**d**,**g**,**j**, From FISH-IF imaging, single plane confocal cross-sections show nuclei (Hoescht, magenta) in wild-type (WT, OreR, **a**), *sry-α -/-* (**d**), *sry-α -/-* plus *sry-α_WT* (**g**), and *sry-α -/-* plus *sry-α_SL1/2^Pt6^* (**j**) embryos. Furrow staining (Peanut, green) allowed measurement of furrow length which serves as a proxy for time (furrow lengths = 5 or 30 μm correspond to Early or Late cellularization, respectively). Orange lines and arrowheads indicate position of the plasma membrane; blue lines and arrowheads indicate position of the apical tops of nuclei. Scale bars = 10 μm**. b**,**e**,**h**,**k**, Nuclear distances for WT (OreR, gray, **b**), *sry-α -/-* (brown, **e**), *sry-α -/-* plus *sry-α_WT* (green, **h**), and *sry-α -/-* plus *sry-α_SL1/2^Pt6^* (purple, **k**) embryos over the course of cellularization (each data point represents nuclear distance for individual nuclei in n ≧ 18 embryos per genotype). Black line shows linear fit for WT data points. **c**,**f**,**i**,**l**, Residual plots show probability of nuclear distance deviations from average WT expected values for WT (OreR, gray, **c**), *sry-α-/-* (brown, **f**), *sry-α -/-* plus *sry-α_WT* (green, **i**), and *sry-α -/-* plus *sry-α_SL1/2^Pt6^* (purple, **l**) embryos (n ≧ 18 embryos per genotype). Black solid lines show Gaussian fit to guide the eye for WT data points; black dashed lines show peak of distribution at residual position = 0 μm for WT embryos. Solid colored lines show Gaussian fit to guide the eye for the other genotypes as indicated; dashed colored lines show peak of distribution at residual position = −1.517, 0 and −0.917 μm for *sry-α -/-* (**f**), *sry-α -/-* plus *sry-α_WT* (**i**), and *sry-α -/-* plus *sry-α_SL1/2^Pt6^* (**l**) embryos, respectively.

Next, we measured nuclear distances in *sry-α -/-* embryos expressing the *sry-α_WT* transgene that encodes apically localizing *sry-α* mRNA (Fig. 2h,j). The *sry-α_WT* transgene restored the increase in nuclear distances as cellularization progressed in the *sry-α* null background (Fig. 5g,h). Nuclear distances were fitted well by the best-fit line for wild-type (Fig. 5h). Residual analysis showed no difference compared to wild-type embryos (Fig. 5i), demonstrating rescue of the nuclear repositioning phenotype by the localizing *sry-α_WT* mRNA.

In contrast, in *sry-α -/-* embryos expressing the *sry-α_SL1/*2*^Pt6^* mRNA, which fails to localize apically (fig. 2i, j), we did not see rescue of the nuclear repositioning phenotype. Nuclei were positioned near the plasma membrane even at late cellularization in these embryos (Fig. 5j). We measured shorter nuclear distances, with most data points falling below the best-fit line for wild-type embryos (Fig. 5k). The residual analysis for *sry-α -/-* embryos expressing the *sry-α_SL1/*2*^Pt6^* mRNA was similar to *sry-α -/-* embryos alone in that the peak probability was shifted below zero towards shorter nuclear distances (Fig. 5l). Therefore, *sry-α* mRNA that fails to localize apically in late cellularization also fails to rescue the nuclear repositioning phenotype observed in *sry-α* null embryos. Note that these embryos have no furrow phenotypes (Fig. 3d), eliminating the possibility that nuclear repositioning secondarily fails in *sry-α -/-* embryos due to furrow regression. We conclude that the cytoplasmic localization of *sry-α* mRNA to the apical tops of nuclei is functionally important for proper nuclear repositioning by late cellularization.

### Translation of apically localizing *sry-α* mRNA is required for nuclear repositioning

Cytoplasmic localization of mRNAs can be important for localized translation and immediate function of the resulting protein ^3,5–8^. To investigate if the translation of apically localizing *sry-α* mRNA is needed to ensure repositioning of nuclei, we used a translation-blocking morpholino ^65,66^. When *sry-α* morpholino was injected in embryos prior to cellularization, it caused cellularization furrows to regress, as previously seen for *sry-α* null and RNAi embryos (Fig. 3b; Extended Data Fig. 5a) ^44^. A control morpholino showed no cellularization defects (data not shown). This suggests that the *sry-α* morpholino can specifically block *sry-α* translation and protein function. When *sry-α* morpholino was injected at cellularization onset, after furrows had already assembled, furrow regression was not observed (Extended Data Fig. 5b). Instead, in these embryos, *sry-α* morpholino disrupted basally-directed nuclear repositioning compared to a control morpholino (Fig. 6a,b). Nuclear distances measured between the apical tops of nuclei and the vitelline membrane were shorter in embryos injected with *sry-α* morpholino compared to control morpholino (Fig. 6c,d; see Extended Data Figs. 6 and 7 for raw and normalized individual embryo data, Extended Data Video 3 and 4). We also saw nuclei popping up, similar to *sry-α* RNAi embryos (compare Fig. 6e,f with Fig. 4f,g), although the tilting phenotype was less obvious. These results support a model whereby localized translation of *sry-α* mRNA in the apical cytoplasm is necessary for proper nuclear repositioning by late cellularization.

**Fig. 6.**
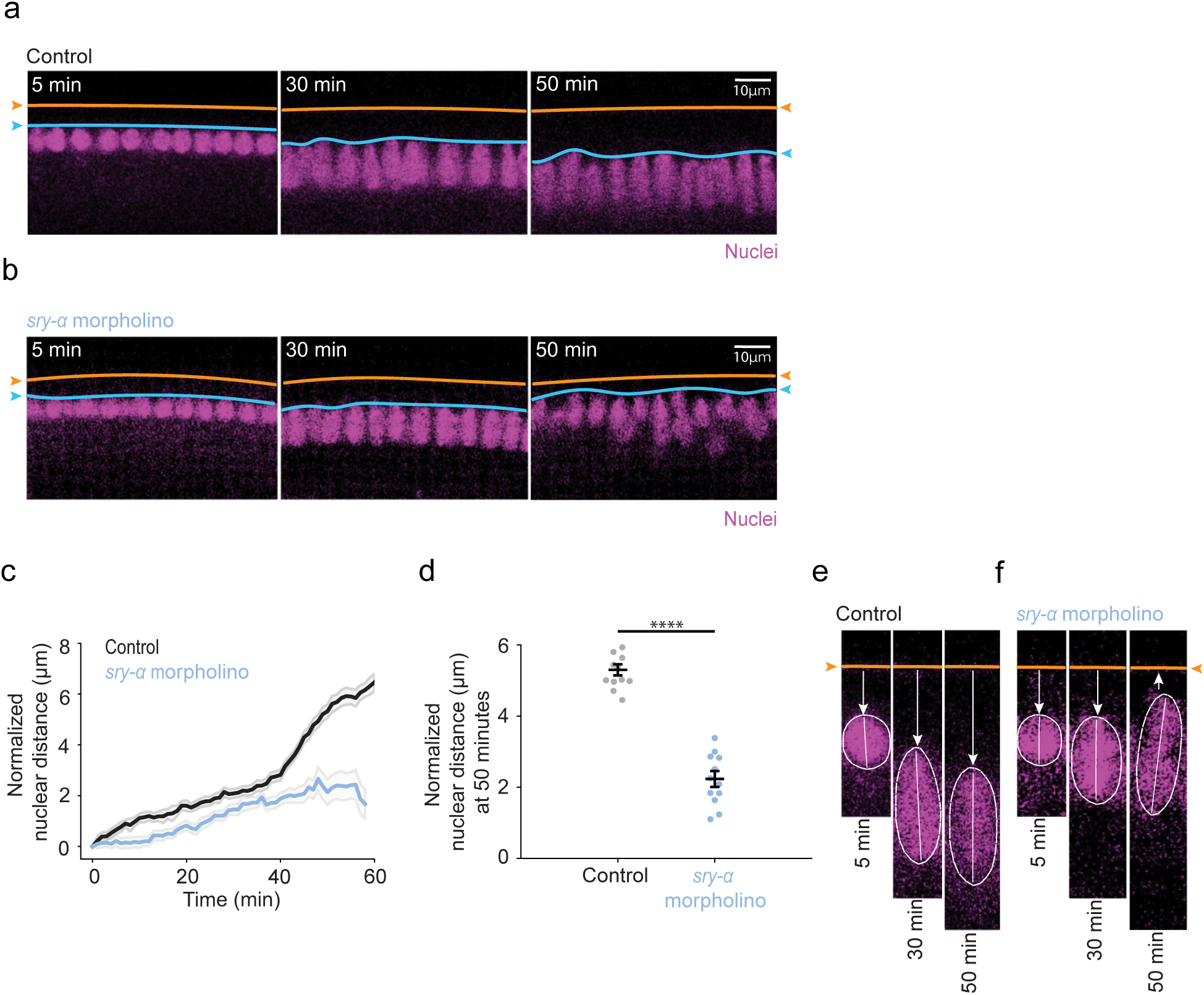
Translation of apically localizing *sry-α* mRNA promotes nuclear repositioning. **a**,**b**, From time-lapse imaging, cross-sections show nuclei (Histone-GFP, magenta) in control morpholino (Control, **a**) or *sry-α* morpholino (*sry-α* morpholino, **b**) injected embryos at 5, 30 and 50 minutes (min) post-cellularization onset. Orange lines and arrowheads indicate position of the vitelline membrane; blue lines and arrowheads indicate position of the apical tops of nuclei. Scale bars = 10 μm**. c**, Normalized nuclear distances for control morpholino (black) and *sry-α* morpholino (blue) injected embryos over the course of cellularization (n = 5 embryos per treatment, ∼15 nuclei followed per embryo; mean ± s.e.m. demarcated in gray). **d**, Normalized nuclear distance at 50 mins post-cellularization onset in control morpholino (gray) and *sry-α* morpholino (blue) injected embryos (each data point represents nuclear distance for ∼15 individual nuclei in n = 5 embryos per treatment). Horizontal lines indicate mean ± s.e.m.; ****p<0.00005, Student’s t test. **e**, **f**, From time-lapse imaging, higher mag cross-sections show an individual nucleus (Histone-GFP, magenta) in control morpholino (Control, **e**) or *sry-α* morpholino (*sry-α* morpholino, **f**) injected embryos at 5, 30 and 50 minutes post-cellularization onset. Orange lines and arrowheads indicate position of the vitelline membrane; white arrows indicate direction of movement of the nucleus; white dashed lines show periphery of the nucleus; white solid lines indicate long axis of the nucleus. **a**,**b**,**e**,**f**, All images are single plane confocal cross-sections.

### Sry-α positively regulates cortical F-actin levels and spot adherens junction assembly

Nuclear positioning frequently depends on the actin cytoskeleton ^23,34–36^. Because Sry-α stabilizes F-actin at furrows in early cellularization ^44,60^, we next asked if it can also regulate the actin cytoskeleton in late cellularization. Based on phalloidin staining for F-actin, we saw a general reduction in cortical F-actin for *sry-α* null versus wild-type embryos even through mid-to-late cellularization (Fig. 7a). We quantified F-actin at the cortex nearby where the *sry-α* mRNA and protein localize in late cellularization and found that F-actin levels in this apicolateral region are significantly reduced in *sry-α -/-* embryos compared to wild-type embryos (Fig. 7a,b). This validates, again, that Sry-α is a positive regulator of F-actin. However, we did not previously recognize that Sry-α’s role in regulating F-actin extended throughout cellularization.

**Fig. 7.**
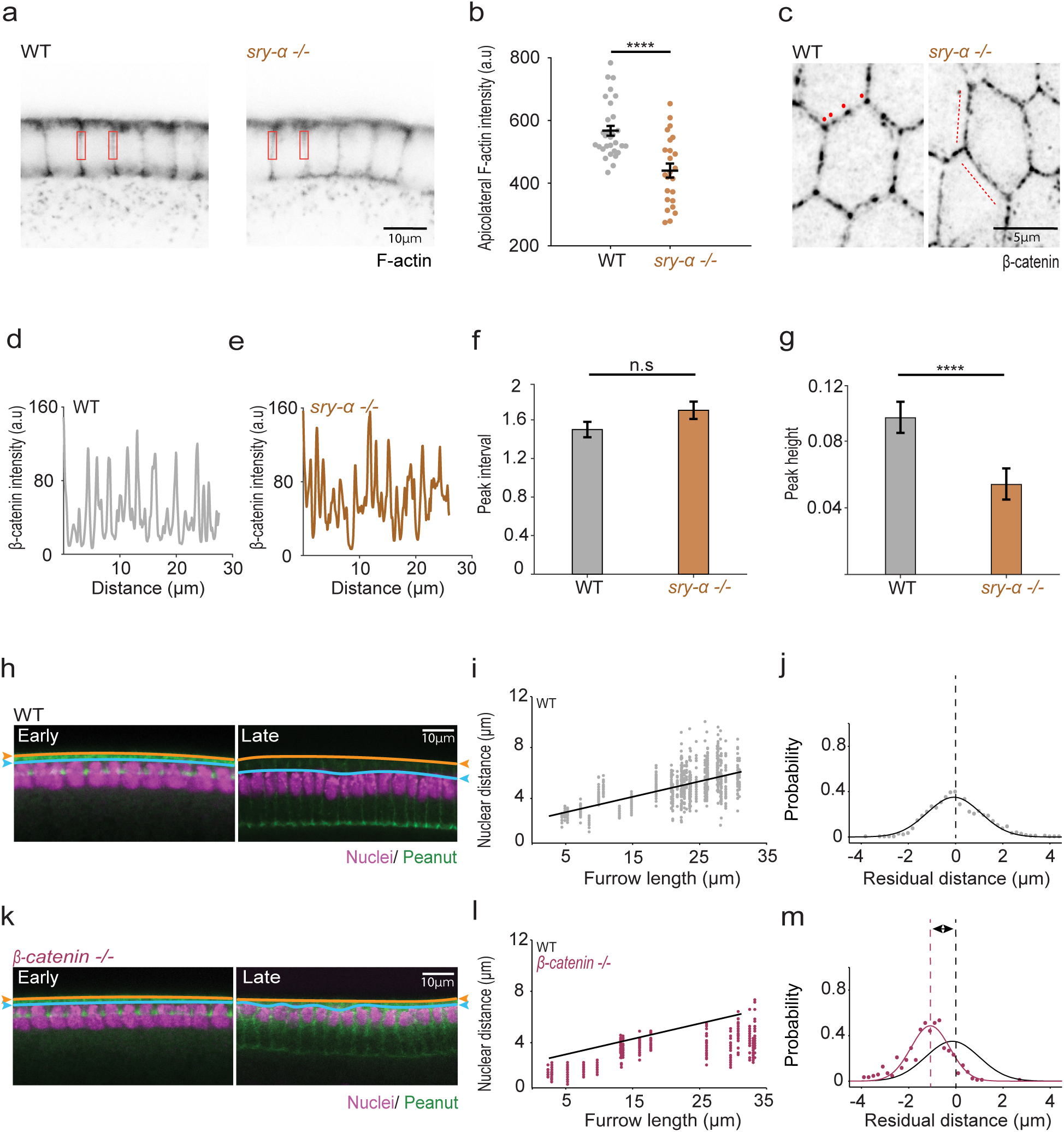
Sry-α promotes assembly of cortical F-actin and spot AJs. **a**, From fixed imaging, inverted cross-sections show cortical F-actin (Phalloidin) in wild-type (WT, OreR, left) and *sry-α -/-* (right) embryos. Red rectangles indicate apicolateral region where F-actin levels were quantified. **b**, Apicolateral F-actin levels in WT (gray) and *sry-α -/-* embryos (brown; each data point represents average intensity for 30 furrows from n = 5 embryos per genotype). Horizontal lines indicate mean ± s.e.m.; ****p< 0.0005, Student’s t test. **c**, From immunofluorescence imaging, inverted XY views show spot AJs (β-catenin) in wild-type (WT, OreR, left) and *sry-α -/-* (right) embryos. Red dots indicate clear periodic puncta in WT embryos; Red dashed lines indicate more evenly distributed signal in *sry-α -/-* embryos. Scale bar = 5 μm. **d**,**e**, Representative β-catenin fluorescence intensity profiles from along the cell-cell interfaces of a representative intact mononucleate cell in wild-type (WT, OreR, **d**) and *sry-α -/-* (**e**) embryos. **f**,**g**, From ACF analysis, peak intervals (**f**) and peak heights (**g**) for WT (gray) and *sry-α -/-* embryos (brown; based on 33 profiles from n = 12 WT or n = 11 *sry-α -/-* embryos; bars indicate mean ± s.e.m.). **h**,**k**, From FISH-IF, cross-sections show nuclei (Hoescht, magenta) in wild-type (WT, OreR, **h**) and *β-catenin -/-* (**k**) embryos. Furrow staining (Peanut, green) allowed measurement of furrow length which serves as a proxy for time (furrow lengths = 5 or 30 μm correspond to Early or Late cellularization, respectively). Orange lines and arrowheads indicate position of the plasma membrane; blue lines and arrowheads indicate position of the apical tops of nuclei. **i**,**l**, Nuclear distances for WT (OreR, gray, **i**) and *β-catenin -/-* (magenta, **l**) embryos over the course of cellularization (each data point represents nuclear distance for individual nuclei in n ≧ 16 embryos per genotype). Black line shows linear fit for WT data points. **j**,**m**, Residual plots show probability of nuclear distance deviations from average WT expected values for WT (OreR, gray, **j**) and and *β-catenin -/-* (magenta, **l**) embryos (n ≧ 16 embryos per genotype). Black solid lines show Gaussian fit to guide the eye for WT data points; black dashed lines show peak of distribution at residual position = 0 μm for WT embryos. Solid magenta line shows Gaussian fit to guide the eye for *β-catenin -/-* embryos (**m**); dashed magenta line shows peak of distribution at residual position = −1 μm for *β-catenin -/-* embryos (**m**). **a**,**c**,**h**,**k**, All images are single plane confocal images. **a**,**h**,**k**, Scale bars = 10 μm. **b**, **f**, **g**, p > 0.05 not significant (n.s.); ****p<0.0005, Student’s t test.

Robust F-actin is needed in the apicolateral domain for assembly of nascent spot adherens junctions (AJs) in late cellularization ^67–69^. So, we also examined AJs in *sry-α* null embryos by immunofluorescence imaging of cell junction component, β-catenin. In wild-type embryos, β-catenin was observed at cell-cell interfaces and concentrated as periodic puncta at spot AJs (Fig. 7c). Spot AJs were also visible in *sry-α -/-* embryos; however, they were not as distinct and their organization was less regular than in wild-type embryos (Fig. 7c). To evaluate whether a difference existed between wild-type and mutant, we prepared fluorescence intensity profiles along cell-cell interfaces, where AJs appeared as peaks of signal (Fig. 7d,e). For *sry-α -/-* embryos, only intact mononucleate cells were analyzed. To assess the spatial organization of junctions, the autocorrelation function (ACF) was calculated from the intensity profiles. Using this analysis, wild-type embryos showed a regular periodicity between junctions, with intervals of 1.5 ± 0.1 μm (mean ± s.e.m.; Fig. 7f). This interval was roughly the same in *sry-α -/-* embryos (1.7 ± 0.1 μm, mean ± s.e.m.; Fig. 7f). However, the peak heights in the ACFs, which provide a measure of the similarity of interval sizes between AJs, showed that *sry-α -/-* embryos are significantly less likely to exhibit the regular periodicity displayed by wild-type embryos (Fig. 7g). This result suggests some perturbation in AJ assembly and/or maintenance in *sry-α -/-* embryos. Thus, both F-actin and junction integrity is compromised in the apicolateral domain of *sry-α* null embryos, nearby the cytoplasmic site where the *sry-α* mRNA localizes.

### AJ assembly is required for nuclear repositioning

Since AJs integrate cytoskeletal systems and support cell structure ^38,39,70^, we reasoned that they might somehow provide important anchor points for nuclei in embryos at late cellularization. If correct, the compromised junctions in *sry-α -/-* embryos might explain why nuclear repositioning fails in these mutants in late cellularization. To test our idea, we prepared embryos maternally deficient in *β-catenin* (*β-catenin -/-*) and assayed basally directed displacement of nuclei. These embryos do not assemble AJs ^71^ and nuclear repositioning failed in these *β-catenin -/-* embryos (Fig. 7h,k), with shorter nuclear distances between the tops of the nuclei and plasma membrane compared to wild-type embryos (Fig. 7i,l). For the *β-catenin -/-* embryos, most data points fall below the best-fit line for wild-type (Fig. 7l). Residual analysis also showed that the peak probability was shifted below zero towards a higher frequency of shorter nuclear distances for *β-catenin -/-* embryos (Fig. 7j,m). These results suggest that AJs contribute to the proper repositioning of nuclei by late cellularization, and we suggest that failed repositioning in *sry-α -/-* embryos could be caused by compromised AJ assembly or maintenance.

## Discussion

Many mRNAs localize to specific subcellular sites, but the functional significance of this localization is frequently unknown. Here, we show that apical localization of *sry-α* mRNA during mid-to-late cellularization promotes polarization of the newly forming cells along their apical/basal axis by enabling spot AJ assembly and basally-directed nuclear repositioning. While it was previously shown that the F-actin binding protein Sry-α acts during early cellularization to stabilize F-actin at ingressing furrow tips ^44,59,60^, a later function remained unrecognized until now. Interestingly, the early function of Sry-α does not require any specific localization of the encoding mRNA, whereas its later activity requires Dynein-mediated accumulation of the mRNA nearby the apical centrosomes. Based on our combined data, we propose that apical *sry-α* mRNA is locally translated to stabilize apicolateral F-actin which supports the assembly of AJs. These AJs then somehow serve as anchor points to ensure proper nuclear repositioning in late cellularization. Additional players are likely to serve as intermediaries between Sry-α, AJs and nuclear repositioning, including microtubules whose disruption perturbs nuclear position in late cellularization ^62^.

Apical localization of *sry-α* mRNA coincides with the establishment of apical/basal polarity in the nascent primary epithelium of the *Drosophila* embryo ^72^. In late cellularization the polarity protein Bazooka/Par3 accumulates as clusters on furrows at the same apical/basal plane as the centrosomes and *sry-α* mRNA localization ^67,68^. These Bazooka clusters then gather intercellular Cadherin/Catenin complexes into spot AJs, which later mature into belt AJs during gastrulation ^73,74^. Bazooka polarization is at first a product of Dynein-based transport and cortical F-actin interaction, and then becomes increasingly refined by mutual exclusion with more apically and more laterally localized polarity proteins, including Par6 and Par1, respectively ^67,68,75–77^. It is possible that locally translated Sry-α promotes initial Bazooka polarization and subsequent spot AJs assembly by reinforcing cortical F-actin next to the centrosomes. This would align with an emerging theme that apical/basal determinants and even AJ components are encoded by mRNAs that localize to the specific region corresponding to their activity along the apical/basal axis ^78–84^.

The distinct subcellular localization of mRNAs has been previously shown to regulate the actin cytoskeleton at sites of dynamic F-actin remodeling. For example, β-actin mRNA concentrates in the axonal growth cones of neurons to support pathfinding and synaptogenesis ^85^, at the leading edge of fibroblasts to promote lamellipodial protrusion and motility ^86^, and at cell-cell and cell-ECM junctions to support cellular adhesion ^82^. Actin nucleators, including the Arp2/3 complex components and Diaphanous/Formin are also encoded by mRNAs that localize to subcellular sites of F-actin dependent processes ^87,88^. In the case of Sry-α, the mRNA localization next to the centrosomes is in proximity to, but not immediately at the apicolateral cell cortex, where F-actin and AJs assemble. While somewhat counterintuitive, this localization is consistent with prior demonstrations that F-actin levels at cellularization furrows depend on activities at this centrosomal position because membrane compartments, including the recycling endosomes, are there and serve as hubs for vesicle transport ^89–91^. Specifically, trafficking through these centrosome-proximal compartments positively regulates the recruitment of RhoGEF2 to the furrow, which activates Rho1, Diaphanous/Formin and cortical F-actin assembly ^89,92,93^. We do not fully understand the molecular details, but vesicles trafficking through these compartments could encounter newly translated Sry-α on their way to the furrow cortex.

From our live data we see that the process of repositioning nuclei during late cellularization is dynamic and requires a balance between opposing forces, as reducing *sry-α* mRNA or blocking its local translation leads to nuclei starting on a basally directed trajectory, but then abruptly reversing to a more apical position. In this study, we have not demonstrated the source of these forces. Active, opposing pushing and pulling forces in the embryo could be exerted on nuclei by F-actin or microtubule -based processes or a combination of both ^33,62,94–98^. While actomyosin-dependent cytoplasmic flows may sweep the nuclei into position ^33,94,97,99^, it is also likely that forces are transmitted to the nuclei via direct cytoskeletal interactions with the LINC complexes, i.e. KASH and SUN proteins that span the nuclear envelope from exterior to interior, respectively ^18,35,37^. In turn, LINC complex components could foster structural integration between nuclei and spot AJs, as LINC components, such as SUN2, have a bidirectional, yet poorly understood, crosstalk with Cadherin-mediated adhesions ^39,100,101^, and we see that AJ perturbation coincides with nuclear mispositioning. Ultimately, the opposing forces arrange nuclei at an initial, single, apical/basal plane throughout the embryo, setting the stage for subsequent, tissue-specific, nuclear migrations that are essential for robust morphogenesis ^63,94,97^. Whatever pushing, pulling or even anchoring mechanisms are at play, our data reveals that mRNA localization serves as an upstream regulator of the force balance required to achieve final apical/basal position of nuclei in the cellularizing embryo.

## Supporting information

Video file related to figure 6

Video file related to figure 6

Video file related to figure 4

Video file related to figure 4

All extended data referenced in the main manuscript

## Acknowledgments

We thank Eric Wieschaus (Princeton University) and Adam Martin (Massachusetts Institute of Technology) for providing fly stocks; Koen Venken (Baylor College of Medicine) for providing constructs and consultation for recombineering, Jordan Raff (University of Oxford) for providing Asterless antibody, Shrunali Amin for assisting with the Ciliobrevin D experiments, and Cuong Diep (Indiana University of Pennsylvania) for feedback and edits. Work in the Sokac lab was supported by the National Institutes of Health (NIH) grants R35 GM136384 and R01 GM115111. Work in the Golding lab was supported by NIH grant R35 GM140709, the National Science Foundation grant 2243257 (NSF Science and Technology Center for Quantitative Cell Biology), and by the Alfred P. Sloan Foundation grant G-2023-19649. Fly stocks were obtained from the Bloomington *Drosophila* Stock Center (supported by NIH grant P40OD018537 and housed at Indiana University – Bloomington). Antibodies against BicD, Septin/Peanut, β-Catenin/Armadillo and Sry-α were obtained from the Developmental Studies Hybridoma Bank (created by the NIH - National Institute of Child Health and Human Development and housed at the University of Iowa – Iowa City).

## Methods

### *Drosophila* stocks and genetics

*Drosophila melanogaster* stocks were maintained at 22°C on standard molasses food. Unless otherwise noted, experiments were performed on embryos in Bownes Stages 4 and 5 ^102^. Oregon R (OreR) was used as the wild-type stock for all experiments as indicated. For rescue experiments, the embryos with one or two copies of endogenous *sry-α* served as wild-type and were identified by RNA-FISH. For imaging *sry-α* mRNA_MS2 in live embryos, F1 embryos were collected from *yw; Histone-RFP; nosMCP-GFP* ^103^ virgin females crossed to *sry-α_MS2* males (generated in this paper; see details below). For *sry-α* RNAi and morpholino injections, embryos were collected from stock *Histone-GFP; Gap43-mCherry*, generated by crossing the original stocks *ubi::H2A-GFP* and *sqh::membrane-mCherry* (provided by Eric Wieschaus, Princeton University, and Adam Martin, Massachusetts Institute of Technology, respectively). For in vitro transcribed mRNA injections, embryos were collected from *ubi::H2A-GFP*. For nuclear distance measurements in fixed embryos, embryos were collected from the following stocks: *sry-α - /TM3Sb, hb::LacZ*, or *sry-α_WT*; *sry-α - /TM3Sb, hb::LacZ*, or *sry-α_SL1/2^Pt6^; sry-α - /TM3Sb, hb::LacZ* (all generated in this paper). For experiments involving *β-catenin -/-* embryos, *β-catenin/armadillo* germ line clones were generated using *arm^2^ FRT 101/FM7a* (Bloomington *Drosophila* Stock Center (BDSC) #8554, Bloomington, IN) and *w* ovo^D1^ v*^24^ *FRT 101/C(1)DX, y^1^ f^1^; hsFLP* (BDSC #1813) stocks.

### Generation of *sry-α _MS2* transgenic stock

To generate the *sry-α_MS2* stock, we used recombineering to insert MS2 stem loops immediately downstream of the stop codon of *sry-α*. A recombineering insert, MS2(24) _Kan, was made in two steps: First, we cloned a series of 24 MS2 stem loops (ACATGGGTGATCCTCATGT) separated by random linker sequences, and a Kanamycin resistance cassette flanked by LoxP sites, into a pBS vector. Second, this vector was used as template to PCR amplify MS2(24)_Kan using primers containing flanking 50 base pair “homology arms” against the *sry-α* target site. We then electroporated the resulting PCR product into SW106 cells (gift of Koen Venken, Baylor College of Medicine) containing the BAC, Ch322-41A22 (BacPac Resources) ^104^. SW106 cells were grown on LB plates with Chloramphenicol and Kanamycin to select for positive recombinants. 20% L-arabinose was used to induce Cre recombinase and excise the Kanamycin cassette in positive recombinants. Cells were then diluted and plated on LB with Chloramphenicol. Ten colonies were selected and replica-plated on both LB-Chloramphenicol and LB-Chloramphenicol/Kanamycin plates to confirm removal of the Kanamycin resistance cassette. The recombineered BAC was isolated and electroporated into EPI300 cells (Epicentre, Madison, WI). Colony PCR, diagnostic restriction enzyme digests, and PCR amplification of the entire *sry-α* region, followed by sequencing, were used to validate the final BAC. For transgenesis, the validated BAC was injected into embryos from stock *y^1^w^67c23^; P{CaryP}attP40* for PhiC31-mediated integration at the attP40 landing site (BestGene, Inc, Chino Hills, CA).

### Generation of *sry-α -/-* stock

To generate *sry-α -/-* flies, we used the CRISPR/Cas system to delete the coding region of the *sry-α* gene ^105^. Two guide RNAs (5’-TATATAGGAAAGATATCCGGGTGAACTTC-3’ and 5’-GACGTTAAATTGAAAATAGGTC-3’) were cloned into vector pCFD4-U6:1_U6:3tandemgRNAs (Addgene plasmid #49411, Watertown, MA). For transgenesis, this vector was injected into embryos from stock *y^1^w^67c23^; P{CaryP}attP2* (BDSC #8622) for PhiC31-mediated integration at the attP2 landing site (BestGene, Inc). Transformants were crossed with stock *y^1^w*nanosCas9* (BDSC #54591) and deletion mutants detected by screening for the known multinucleation phenotype of *sry-α* null mutants ^44,47,59,60^. To validate the *sry-α* deletion, rescue was confirmed with a *sry-α* genomic rescue stock (gift of Eric Wieschaus) ^44^. For experiments, the deletion was balanced to make *sry-α - /TM3Sb, hb::LacZ*.

### Synthesis of *in vitro* transcribed fluorescent mRNAs

Constructs for *in vitro* transcription were generated by cloning each gene or gene fragment of interest in a pCS2+ vector. For *sry-α, bicoid, and string* full-length constructs, full-length gene sequences were amplified by PCR from cDNAs, then cloned by restriction enzyme digestion and ligation downstream of the SP6 promoter in pCS2+. All deletion and 3’UTR replacement constructs were derived from these full-length constructs using PCR-based strategies. To generate *sry-α_SL1/2^Pt3^ and sry-α_SL1/2^Pt6^*, two DNA sequences consisting of the *sry-α* 3’ UTR with mutations in three or six nucleotides involved in stem loop base-pairing, respectively, were commercially synthesized (GeneWiz, South Plainfield, NJ) and cloned into pCS2+_*sry-α_ΔUTR* by restriction enzyme digestion and ligation. For fluorescent mRNA synthesis, these pCS2+ constructs were linearized and used as a template for *in vitro* transcription (mMessage mMachine SP6 kit, Thermo Fisher Scientific, Waltham, MA), with 5nmol Cy5-UTP (Perkin Elmer, Waltham, MA) added per 10μL reaction. TURBO DNase (Thermo Fisher Scientific) was used to eliminate the template, followed by Lithium Chloride precipitation, per kit instructions. RNA was diluted to a working concentration of 1mg/ml.

### RNA secondary structure prediction

To predict secondary structures in the *sry-α* mRNA and select mutations to disrupt the stem loops in the 3’ UTR, the *sry-α* mRNA sequence was analyzed using RNAstructure (https://rna.urmc.rochester.edu/RNAstructureWeb/Servers/Predict1/Predict1.html) ^106^, with default settings.

### Generation of sry-α_WT, sry-α_SL1/2^Pt3^, and sry-α_SL1/2^Pt6^ stocks

To generate *sry-α_WT, sry-α_SL1/2^Pt3^,* and *sry-α_SL1/2^Pt6^* stocks, PCR was used to amplify the *sry-α_WT, sry-α_SL1/2^Pt3^,* and *sry-α_SL1/2^Pt6^* sequences previously cloned into pCS2+ vectors, with accompanying *β-globin* promoter/5’UTR sequences. PCR products were inserted into attB-Pacman-Amp^R^ (gift of Koen Venken) by restriction enzyme digestion and ligation, and constructs were transformed into Epi300 cells (gift of Koen Venken). Insertion of the correct sequences was confirmed using PCR amplification and sequencing of the full gene region. For transgenesis, constructs were injected into embryos from stock *y^1^w^67c23^; P{CaryP}attP40* for PhiC31-mediated integration at the attP40 landing site (BestGene, Inc). For experiments that required transgenic constructs to be expressed in a *sry-α -/-* background, transgenic flies were crossed with *sry-α -/TM3Sb, hb::LacZ* so that endogenous *sry-α -/-* could be detected by RNA-FISH, looking for absence of *LacZ mRNA*.

### Embryo collection and fixations

Embryo collection cups were set up at room temperature, unless otherwise stated, on yeasted apple juice agar plates according to published protocols ^107^. Embryos were dechorionated in bleach for 1 minute. For RNA-FISH, embryos were fixed at the interface of 4% paraformaldehyde in PBS and n-heptane (1:1) for 15 minutes and released from the vitelline membrane by methanol popping. For immunofluorescence, embryos were fixed at the interface of 4% paraformaldehyde in 0.1M phosphate buffer (pH 7.4) and n-heptane (1:1) for 15 minutes and released from the vitelline membrane by methanol popping. For nuclear distance measurements using FISH-IF, embryos were fixed at the interface of 8% paraformaldehyde in PBS and n-heptane (1:1) for 30 minutes and released from the vitelline membrane by hand peeling. For phalloidin staining, embryos were fixed in 16% PFA in 1x PBS and n-heptane (1:1) for 20 minutes and released from the vitelline membrane by hand peeling. For EPS permeabilization and Ciliobrevin D treatment, embryos were fixed at the interface of 4% paraformaldehyde in PBS and n-heptane (1:1) for 15 minutes and released from the vitelline membrane by methanol popping.

### Antibodies

The following antibodies were used for immunofluorescence or FISH-IF: Peanut/Septin (1:500 dilution, 4C9H4, Developmental Studies Hybridoma Bank (DSHB), Iowa City, Iowa); Asterless (1:500 dilution, gift from Jordan Raff, University of Oxford); β-Catenin/Armadillo (1:50 dilution, N27A1, DSHB); Sry-α (1:200 dilution, 1G10, DSHB, concentrated using ammonium sulfate precipitation). The following antibodies were used for immunoblots: Sry-α (1:200 dilution, 1G10, DSHB, concentrated using ammonium sulfate precipitation); β-actin C4 (1:200 dilution, sc-47778, Santa Cruz Biotechnology, Dallas, TX). The following antibodies were used for BicD immunoprecipitation: Bicaudal-D (1B11 and 4C2, DSHB, combined in 1:1 ratio for a total 12µg antibody per pull-down).

### RNA-FISH, FISH-IF and immunofluorescence

For RNA-FISH, the protocol was adapted from ^108^. FISH probes complementary to target mRNAs (72 probes for *hbLacZ* and 66 probes for *sry-α*, respectively; Stellaris, Inc, Petaluma, CA) were 3’ TAMRA labeled ^108^. Fixed embryos were rehydrated by washing 4 x 10 minutes in PBTx (1x PBS, 0.1% Triton X-100), followed by 2 x 10 minutes in wash buffer (2x SSC, 0.01% Triton X-100, 20% formamide), and then hybridized with 500nM FISH probes in hybridization buffer (10% Dextran sulfate, 0.1% E.coli tRNA, 2mM Vanadyl ribonucleoside, 0.2mg/ml RNAse free BSA, 2x SSC, 20% formamide) overnight at 30°C. Embryos were then subjected to post-hybridization washes 3 x 30 minutes in wash buffer at 30°C, and stained for DNA using Hoechst 33342 (1:1000 dilution, Thermo Fisher Scientific) during the last wash. Embryos were mounted in AquaPolymount (Polysciences, Warrington, PA).

For combined FISH-IF, immunofluorescence was carried out first by rehydrating embryos 4 x 10 minutes in PBTx (1x PBS, 0.1% Triton X-100) and then blocking with PBTB (1x PBS, 20% Western Blocking reagent, 2mM Vanadyl ribonucleoside, 0.01% Triton X-100) for 1 hour at room temperature. Embryos were washed 4 x 10 minute with PBTx and incubated with primary antibody for 1 hour at room temperature. This was followed by washing 4 x 10 minute in PBTx and incubation with secondary antibody for 1 hour at room temperature. Embryos then underwent washing 4 x 10 minute in PBTx followed by 2 x 5 minute with wash buffer (2x SSC, 0.01% Triton X-100, 20% formamide). Embryos were hybridized with 500nM FISH probes in hybridization buffer (10% Dextran sulfate, 0.1% *E. coli* tRNA, 2mM Vanadyl ribonucleoside, 0.2mg/ml RNAse free BSA, 2x SSC, 20% formamide) overnight at 27°C and washed 2 x 30 minutes with wash buffer at 27°C. DNA was stained using Hoechst 33342 (1:1000 dilution) in the last wash. Finally, embryos were washed 2 x 10 minutes in 2xSSC + 0.1% Triton X-100 and mounted in Prolong Diamond Antifade Mountant (Thermo Fischer Scientific).

For immunofluorescence, embryos were rehydrated 4 x 10 minutes in 1% BSA in PBTx (1x PBS, 0.1% Triton X-100), followed by blocking in 10% BSA in PBTx for 1 hour at room temperature. Embryos were incubated with primary antibody in 5% BSA PBTx for either 1 hour at room temperature or overnight at 4°C. Embryos were washed 4 x 10 minutes in 1% BSA PBTx and then incubated with secondary antibody in 5% BSA PBTx for 1 hour at room temperature. After this, embryos were washed 1 x 10 minutes in 1% BSA PBTx and stained for DNA using Hoechst 33342 (1:1000 dilution) in 1% BSA PBTx for 10 minutes. Finally, embryos were washed 4 x 10 minutes in 1% BSA PBTx and then mounted on slides using Prolong Diamond Antifade Mountant.

### BicD immunoprecipitation and QPCR

BicD immunoprecipitation was performed as in Vazquez-Pianzola et al., 2017 ^52^ with the following modifications. An aliquot of 50µl Dynabeads (Invitrogen, Carlsbad CA) was briefly washed with 1x PBST (1x PBS, 0.1% Tween-20), incubated with 12µg BicD antibody for 20 minutes at room temperature, and then washed with non-hypotonic buffer (20mM HEPES pH 7.9, 2mM MgCl_2_, 150 mM KCL, 1mM DTT, 20% glycerol, 0.5% Tween-20, 1X EDTA-free protease inhibitor (Roche, Indianapolis, IN)). Negative control beads were treated the same but without antibody.

For lysate preparation, 400 embryos were handpicked at mid-cellularization and lysed at 4°C in 800µl hypotonic buffer (20mM HEPES pH 7.9, 2mM MgCl2, 10mM KCL, 1mM DTT, 0.5% Tween-20, 1X EDTA-free protease inhibitor (Roche), 1 unit/ul RNase inhibitor (New England Biolabs, Ipswich MA)). Lysate was centrifuged at 14,000 g for 10 minutes and 600µl of the clear middle layer was collected and combined with 323µl high salt buffer (20mM HEPES pH 7.9, 0.4M KCL, 2mM MgCl_2_, 1mM DTT, 57% glycerol, 2U/100µl RNAse-free DNAse 1, New England Biolabs, Ipswich, MA). This mixture was precleared, divided equally between the control and BicD antibody coated beads, and incubated overnight at 4°C. Collected beads were washed 8 x 10 minutes with high salt buffer and treated with 30µg Proteinase K (Invitrogen) in 100µl protease buffer (150 mM NaCl, 12.5 mM EDTA, 10% SDS, 0.1M Tris-HCl pH 7.5) for 30 minutes at 55°C. To extract RNA, TRIzol (Thermo Fisher Scientific) was added according to manufacturer’s instructions with 5 µg RNA grade glycogen included to improve RNA recovery. cDNA was synthesized using iScript Select cDNA Synthesis Kit (Bio-Rad, Hercules, CA). QPCR was performed using the PowerUp SYBR Green Master Mix (Applied Biosystems, Waltham, MA) on the StepOnePlus System (Applied Biosystems). Fold change was calculated using the ΔΔCT method ^109^ and fold change in the BicD versus no antibody control were plotted (n=3 independent biological replicates).

**Table 1.**
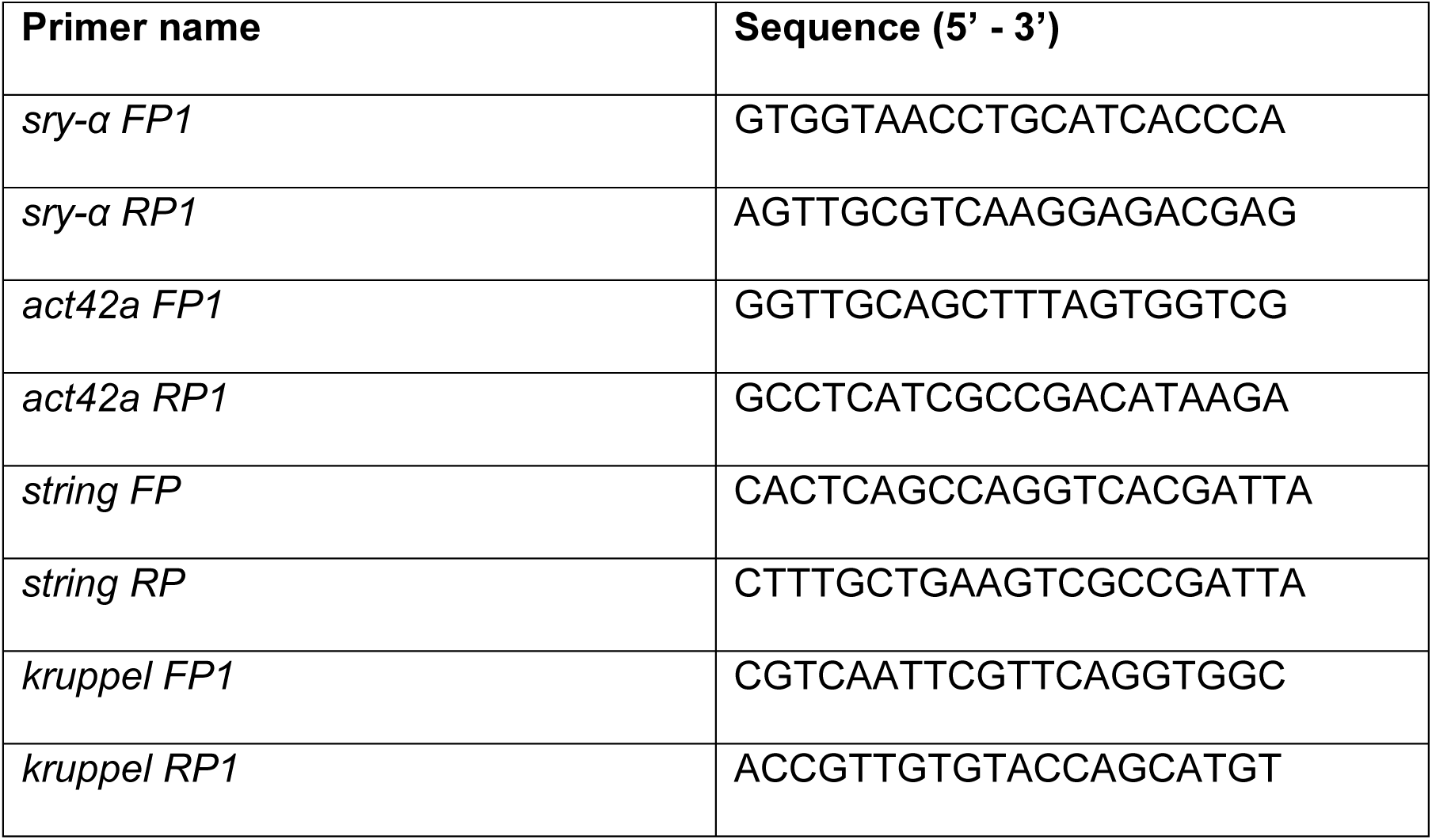
Primers used for QPCR following BicD immunoprecipitation.

### Embryo injections

For injection of *in vitro* transcribed mRNAs, *ubi::H2A-GFP* embryos were dechorionated in bleach for 1 minute, mounted on a cover slip using “embryo glue”, dehydrated, and covered with a 1:3 mix of Halocarbon 700: Halocarbon 27 oil (Millipore Sigma, Burlington, MA). Embryos at early cellularization were identified by eye under a dissecting microscope, then injected with ∼50 pl of RNA using Eppendorf Femtojet injection system (Eppendorf, Framingham, MA). Embryos were immediately moved to the microscope for time-lapse imaging.

For RNAi, *Histone-GFP; Gap43-mCherry* embryos were processed as described above and then injected with ∼50 pl of *sry-α* specific dsRNA. Embryos were moved to a 25°C incubation chamber for 30 minutes then to the microscope for time-lapse imaging.

For injection of morpholinos, *Histone-GFP; Gap43-mCherry* embryos were processed as described above and then injected with ∼50pl of 0.5μM control or translation blocking morpholino (control sequence 5’-GAAATGTTCAACTAAGTTCTGGCGG-3’; *sry-α* morpholino sequence 5’-TTCCATGCTGTTCTATCAGATG-3’, Gene Tools, LLC Philomath, OR). Morpholinos were injected at either the end of nuclear cycle 13 or beginning of nuclear cycle 14 as indicated. Embryos were immediately moved to the microscope for time-lapse imaging.

### EPS permeabilization and Ciliobrevin D treatment

For permeabilization, EPS solvent (90% d-limonene, 0.5% Cocamide DEA and 0.5% Ethoxylated alcohol) was diluted 1:40 in PBS. Embryos were dechorionated in bleach for 1 minute, immersed in diluted EPS for 90 seconds, washed 6 x 10-15 seconds in PBS, transferred to a glass scintillation vial with 80μM Ciliobrevin D (Millipore Sigma) for 15 minutes and then immediately fixed.

### Microscopy

Images were collected on Zeiss LSM 710 or 880 confocal microscopes using either a 40x water-immersion objective (NA 1.2) or 63x oil-immersion objective (NA 1.4) and a pinhole of 1 AU. For all live imaging, embryos were dechorionated and mounted as previously described ^107^. For *sry-α_MS2* imaging, Z-stacks comprised of 1μm thick slices encompassing ∼12μm of the apical surface were captured at 5-minute intervals, beginning in nuclear cycle 13 and continuing through the end of cellularization. For RNAi and morpholino injected embryos, single-plane cross-sectional images were captured at the embryo equator on the ventral side at 1-minute intervals, beginning at the end of nuclear cycle 13 and continuing through the end of cellularization, extended data video 3 was captured using Airyscan. For *in vitro* transcribed mRNA experiments, single-plane cross-sectional images were captured at the embryo equator at 2-minute intervals for 30 minutes. For nuclear distance experiments with fixed embryos, single-plane cross-sectional images were captured at the embryo equator on the ventral side. For RNA-FISH and FISH-IF for genotyping, Z-stacks comprised of 0.25μm thick slices encompassing ∼7μm were captured at the apical surface of the embryo. For multinucleation experiments, single-plane images were collected at the plane of the furrow tips. For F-actin levels, single-plane cross-sectional images were captured at the embryo equator on the ventral side. For β-Catenin images, Z-stacks comprised of 0.3μm thick slices encompassing 5μm were collected at the apical surface of the embryo using the Zeiss 880 with Airyscan.

### Image processing

All images were analyzed and processed in Image J2/Fiji (https://fiji.sc). Graphs were generated in MATLAB (MathWorks, Natick, MA) and Python (Spyder compiler in Anaconda) and edited in Adobe Illustrator CC (Adobe, San Jose, CA). Figures were prepared in Adobe Photoshop CC.

### Statistical analysis

Student’s t-tests were performed using GraphPad QuickCalcs (GraphPad, San Diego, CA). One-way ANOVA tests were performed using https://statpages.info/anova1sm.html website. P values ≤0.05 were considered statistically significant.

### Densitometry analysis for immunoblots and RT-PCR

Densitometry was carried out in Image J2/Fiji (https://fiji.sc). Gel bands were selected and brightness measured. Following this integrated intensity was calculated by multiplying the brightness with area of box used for selection. Normalization factor was calculated by dividing the wild-type integrated intensity for actin (*actin42a* for RT-PCR or β-actin for immunoblots) by the integrated intensity for the other genotypes. This normalization factor was then used to calculate normalized *sry-α* mRNA or Sry-α protein intensity by multiplying the integrated intensity with the normalization factor.

### Apical localization quantification for *sry-α_MS2* and endogenous *sry-α*

For quantification of MS2 foci in live embryos, time-lapse movies were split into individual time points. The Z-stack image for each time point was used to generate four YZ-projections every 100 pixels (∼14μm) across the long axis of the embryo. This method allowed us to collect information from a sample of ∼50 nuclei per time point per embryo in an unbiased fashion. Visible nuclei, nuclear foci, and apical foci in each YZ projection were then counted and summed for all the YZ projections per time point. The fraction of nuclei containing nuclear foci (nuclear foci / nuclei) and the fraction of nuclei with apical foci (apical foci / nuclei) were then calculated for each time point.

For quantification of apical *sry-α* mRNA after RNA-FISH, images were analyzed by making maximum intensity projections of Z-stacks and cropping the image to 40×40 in the area with maximum signal followed by using the despeckle function and autolocal threshold using the Bernsen method. mRNA spots were measured using the analyze particles function with particle size range 0.15-infinity.

### Apical localization quantification for *in vitro* transcribed mRNAs

For quantification of apical localization of mRNAs in live injected embryos, images were analyzed by measuring the Cy5 fluorescence intensity in fixed-size boxes near the injection site at positions apical and basal to the nuclei immediately post-injection (T0) and again 30 minutes post-injection (T30). The ratio of apical to basal signal was calculated at each time point, and the T30 ratio was subtracted from the T0 ratio to give Δapical: basal^30min^. This quantification was also done using apical and basal signal at 0- and 10-minutes post-injection and gave comparable results.

### Multinucleation quantification

To quantify the penetrance of embryos exhibiting multinucleation, we examined 169 μm^2^ single-plane surface views encompassing Peanut stained furrow tips, which allowed us to view ∼1000 nuclei simultaneously. Multinucleation was treated as binary: embryos with any visible multinucleate cells were counted as “multinucleate” with furrow regression and embryos with no visible multinucleate cells were counted as “not multinucleate” with successful or rescued furrow ingression.

### Nuclear distance quantification

First, furrow lengths were measured using custom MATLAB code. Final average furrow lengths per embryo were calculated based on the average furrow length from 5 furrows on the dorsal side and 5 furrows on the ventral sides of the embryo. Then, nuclear distance was measured by drawing a straight line from the apical top of nuclei that were in focus to either the vitelline membrane (live embryos) or plasma membrane (fixed embryos). For fixed embryos, 20-25 nuclei were quantified on both the dorsal and ventral side per embryo. For live embryos, 12-15 nuclei were quantified per embryo at each time point from the end of nuclear cycle 13 and continuing through the end of cellularization. Raw nuclear distances were normalized by subtracting the distances for every time point from nuclear distance at 0-minute time point. Residual analysis was carried out using custom Python code using the Plotly, Pandas, NumPy and SciPy libraries. Residuals were calculated by fitting the data to a linear model using least squares regression and then subtracting the observed value from predicted value from the linear fit. Residual arrays were created for each genotype and normalized residuals were calculated by dividing the residual value by the predicted value. This allowed for expression of residuals as a percentage error of the predicted value. Histograms were then generated using the normalized residuals showing the distribution of deviations from the fitted model for each genotype. A Gaussian was fitted to each genotype to compare these deviations.

### Apicolateral F-actin quantification

First, furrow lengths were measured using custom MATLAB code. Final average furrow lengths per embryo were calculated based on the average furrow length from 5 furrows on the dorsal side and 5 furrows on the ventral sides of the embryo. Then, apicolateral F-actin intensity was measured for 5 furrows on the ventral side per embryo by drawing a 4.5 μm^2^ box along the length of each furrow starting just below the apical embryo surface. Care was taken not to include F-actin fluorescence from the apical embryo surface. Only intact furrows were used for measurement in *sry-α -/-* embryos. Control and *sry-α -/-* embryos were stained in the same tube to avoid batch effects between tubes.

### AJs quantification and statistical analyses

First, furrow lengths were measured using custom MATLAB code. Final average furrow lengths per embryo were calculated based on the average furrow length from 5 furrows on the dorsal side and 5 furrows on the ventral sides of the embryo. AJ quantification was limited to embryos with furrow lengths between 25 and 35μm, and only intact furrows were used for measurement in *sry-α -/-* embryos. From single-plane surface views, fluorescence intensity profiles were generated by tracing furrows (i.e. cell interfaces) using the segmented line tool in Image J2/Fiji and an equal number of profiles (n=33) were analyzed from 12 OreR embryos and 11 *sry-α -/-* embryos. For each profile, the fluorescence intensity 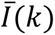 was sampled along its length at a fixed spatial interval of 0.0668 µm. To assess the periodicity of the fluorescence signal, the spatial autocorrelation function (ACF) of each intensity profile was computed using the following formula:

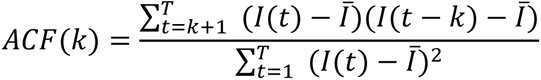

where 𝑘 is the displacement and 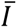 is the mean intensity across the furrow. The ACF was computed up to 𝑘 = 𝑇 = 130. To quantify periodicity, the peak height and peak interval of the ACF were analyzed. Peaks were identified using the scipy.signal.find_peaks function in Python, with a minimum peak distance of 10 sampling units (0.668μm).

**Table 2.** Constructs used for *in vitro* transcription and injection.

| Construct name | Description |
| --- | --- |
| pCS2_*sry-α*_WT | Wild-type, full-length *sry-α* construct |
| pCS2_*sry-α*_ΔUTR | *sry-α* construct lacking the 3' UTR |
| pCS2_*sry-α*_SL1 | *sry-α* construct containing only the first 109 (of 217) nucleotides of the 3' UTR (second stem loop deleted) |
| pCS2_*sry-α*_SL2 | *sry-α* construct containing only the last 98 (of 217) nucleotides of the 3' UTR (first stem loop deleted) |
| pCS2_*sry-α*_SL1/2<sup>Pt3</sup> | *sry-α* construct with three point mutations designed to disrupt both stem loops |
| pCS2_*sry-α*_SL1/2<sup>Pt6</sup> | *sry-α* construct with six point mutations designed to disrupt both stem loops |
| pCS2_*bicoid*_WT | Wild-type, full-length *bicoid* construct (positive control for apical localization) |
| pCS2_*string*_WT | Wild-type, full-length *string* construct (negative control for apical localization) |
| pCS2_*string*_sry-α | *string* construct with its native 3' UTR replaced by the *sry-α* 3' UTR |

---

**Extended Data Fig. 1 | Related to Fig. 3e-g.**
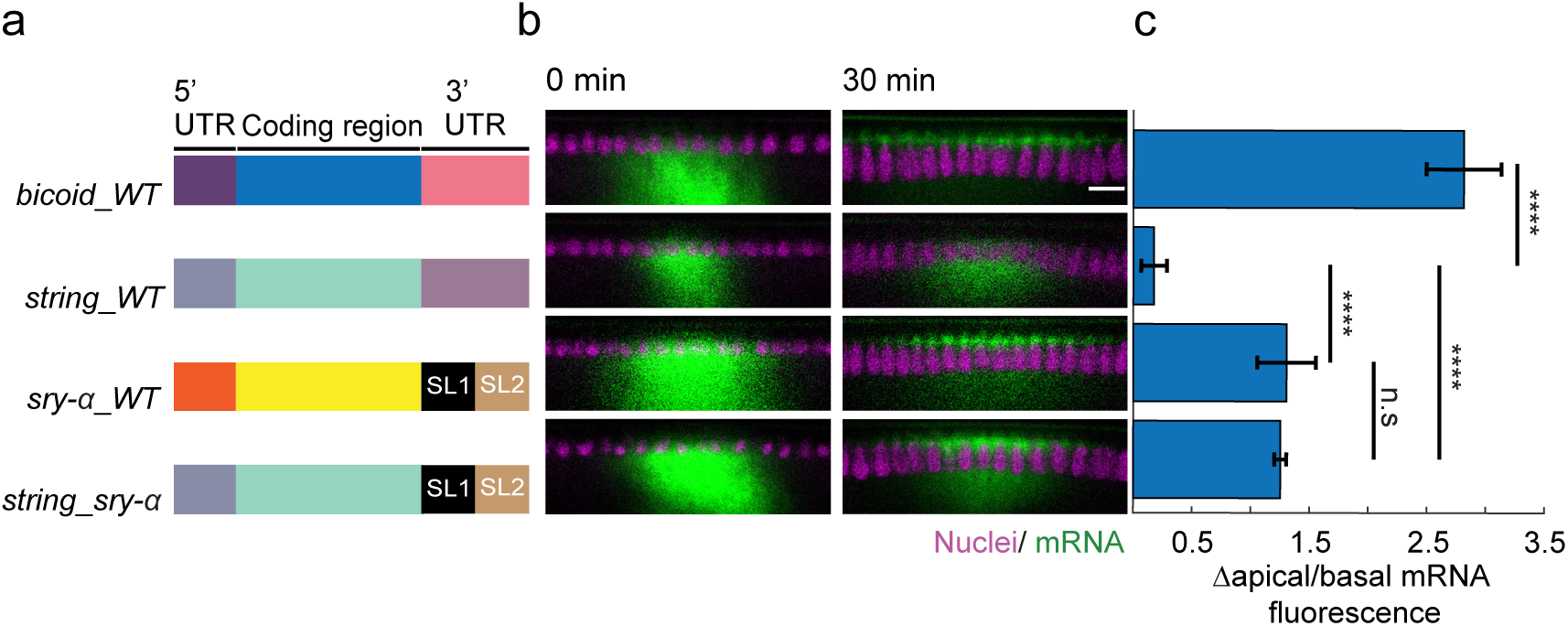
**a**, Schematic shows construct design for *in vitro* transcribed mRNAs for positive control (*bicoid_WT*), negative control (*string_WT*) and *sry-α* experimentals (*sry-α_WT*, *string_sry-α*; WT, wild-type; SL, stem loop). **b**, From time-lapse imaging, cross-sections show injected mRNA (green) and nuclei (Histone-mCherry, magenta). Identity of the mRNA is indicated by aligned construct in (**a**). Imaging started immediately after injection and proceeded for 30 minutes (0 min and 30 min, respectively). Scale bar = 10 μm. **c**, Change in apical to basal fluorescence from 0 to 30 minutes for the mRNA indicated by aligned construct in (**a**). Bars indicate mean ± s.e.m. (n = 5 embryos per construct); p > 0.05, not significant (n.s.); ****p<0.00005, one-way ANOVA.

**Extended Data Fig. 2 | Related to Fig. 3h, i.**
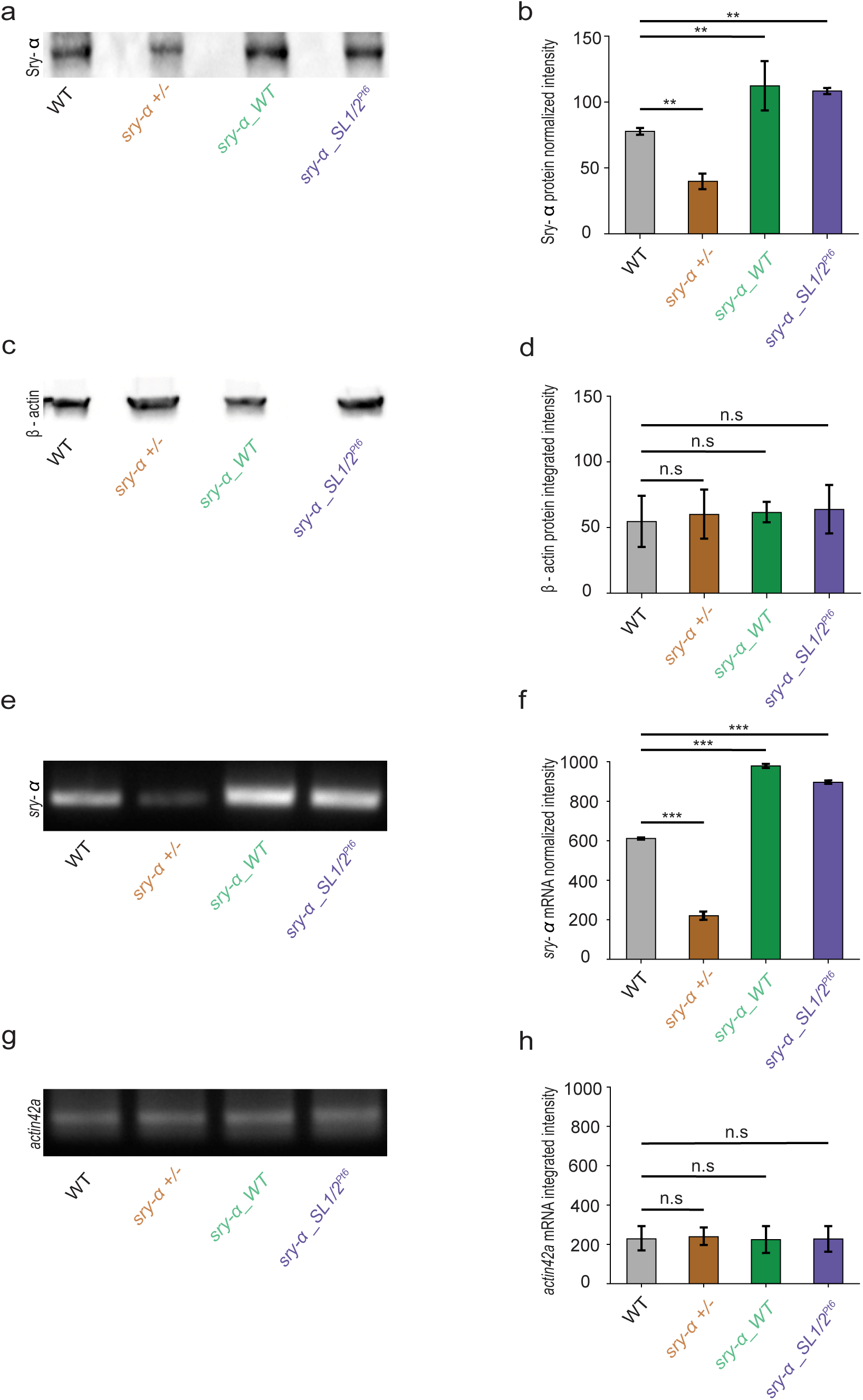
**a**,**c**, Immunoblots show Sry-α and β-actin bands, respectively, for protein lysates from embryos from indicated stocks. **b**,**d**, Quantification of Sry-α and β-actin protein levels, respectively, as determined by densitometry. **e**,**g**, Ethidium bromide stained agarose gels show reverse transcription-PCR products for *sry-α* and *actin42a*, respectively, following mRNA extraction from embryos from indicated stocks. **f**,**h**, Quantification of *sry-α* and *actin42a* PCR product levels, respectively, as determined by densitometry. PCR product levels are considered a proxy for mRNA levels in embryos. **a**-**h**, Stocks are: OreR (wild-type, WT); *sry-α - /TM3Sb, hb::LacZ (sry-α +/-)*; *sry-α_WT; sry-α_SL1/2^Pt6^*. **b**,**d**,**f**,**h**, Bars indicate mean ± s.e.m. (n = 3 biological replicates); p > 0.05, not significant (n.s.); **p<0.005, ***p<0.0005, ****p<0.00005, Students t-test.

**Extended Data Fig. 3 | Related to Fig. 4.**
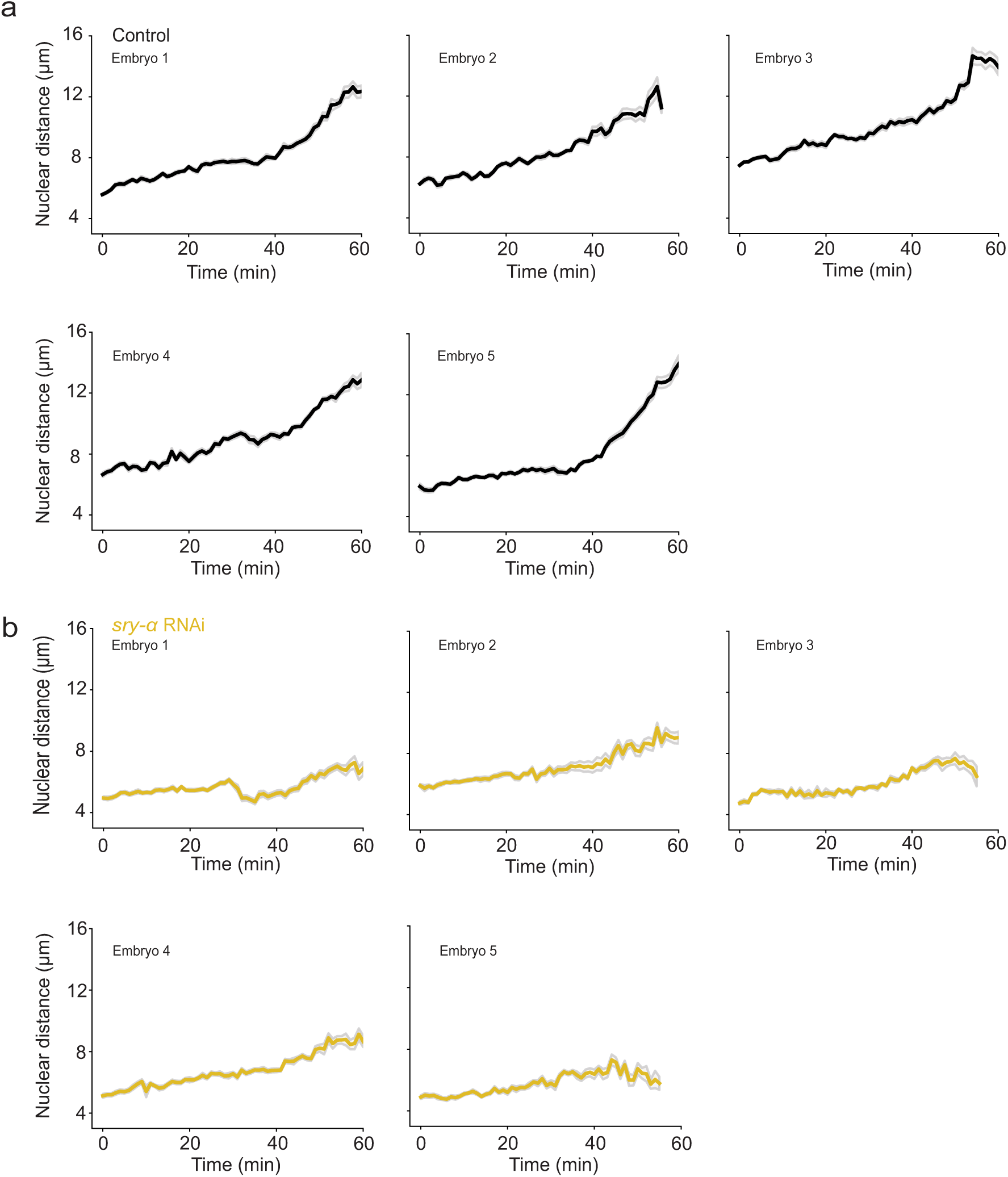
**a**,**b**, Individual embryo, raw nuclear distances for buffer (Control, black, **a**) and *sry-α* dsRNA (*sry-α* RNAi, gold, **b**) injected embryos over the course of cellularization (15 nuclei followed per embryo; mean ± s.e.m. demarcated in gray).

**Extended Data Fig. 4 | Related to Fig. 4.**
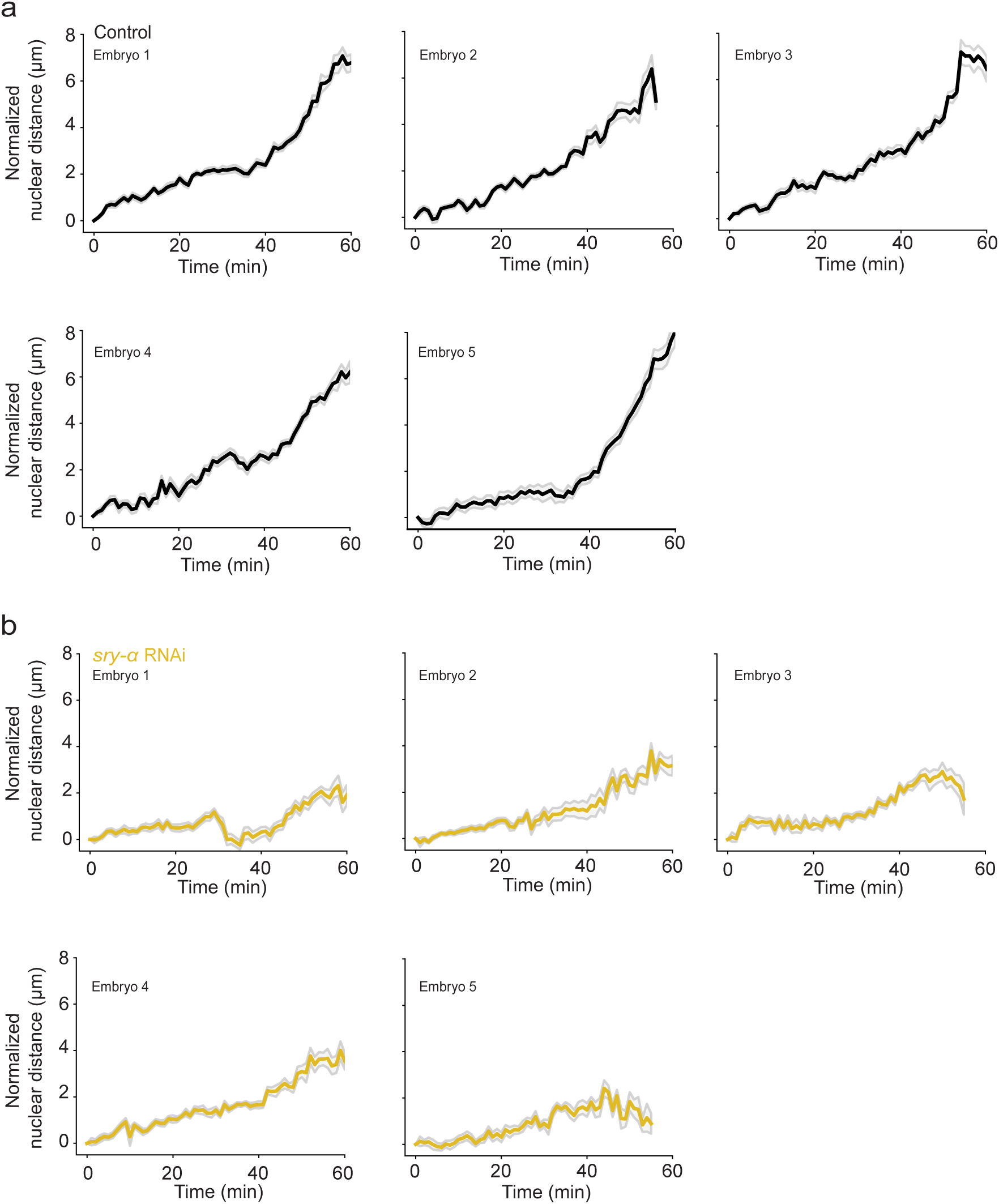
**a**,**b**, Individual embryo, normalized nuclear distances for buffer (Control, black, **a**) and *sry-α* dsRNA (*sry-α* RNAi, gold, **b**) injected embryos over the course of cellularization (15 nuclei followed per embryo; mean ± s.e.m. demarcated in gray).

**Extended Data Fig. 5 | Related to Fig. 6.**
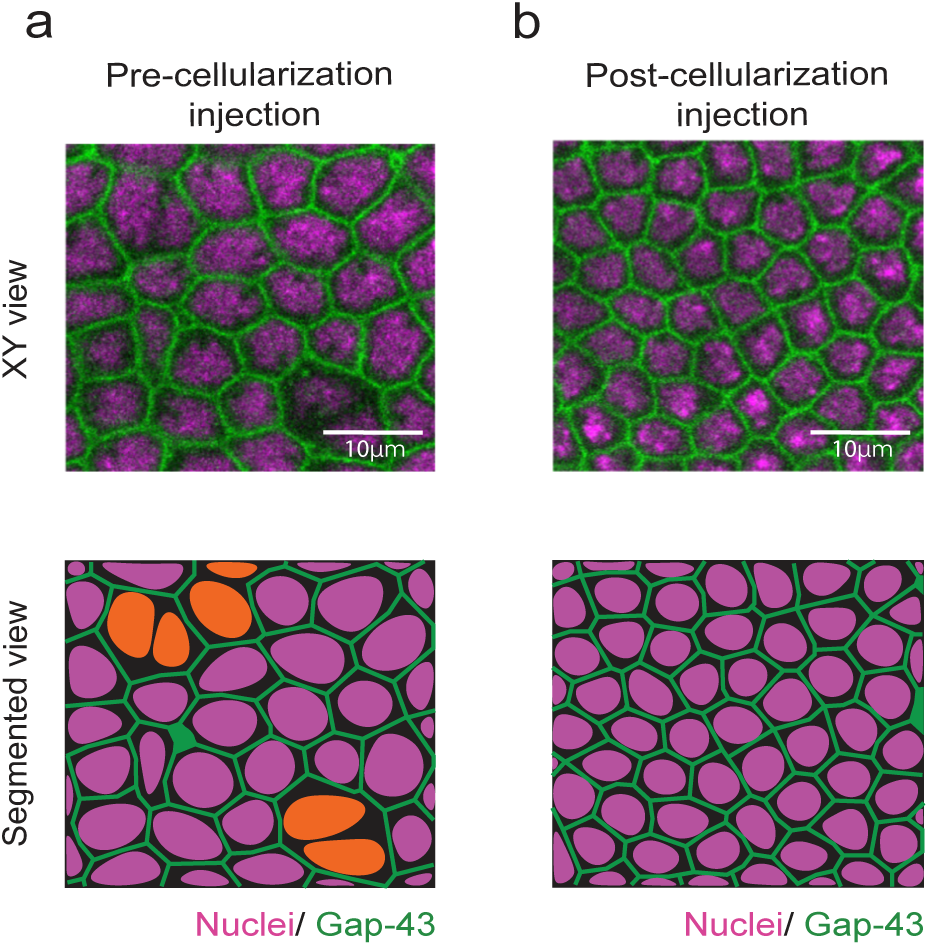
**a**,**b**, From time-lapse imaging, single plane confocal XY views show cellularization furrows (Gap43-GFP / plasma membrane, green) ingressing between nuclei (Histone-mCherry, magenta) in *sry-α* morpholino injected embryos. Embryos were either injected prior to cellularization (**a**) or just after cellularization onset (**b**; n = 5 embryos per treatment). Bottom row shows segmented views of the corresponding XY views with nuclei highlighted (orange, **a**) in multinucleated cells where furrows regressed. Images collected from embryos at late cellularization, with furrow lengths > 5 μm. Scale bars = 10 μm.

**Extended Data Fig. 6 | Related to Fig. 6.**
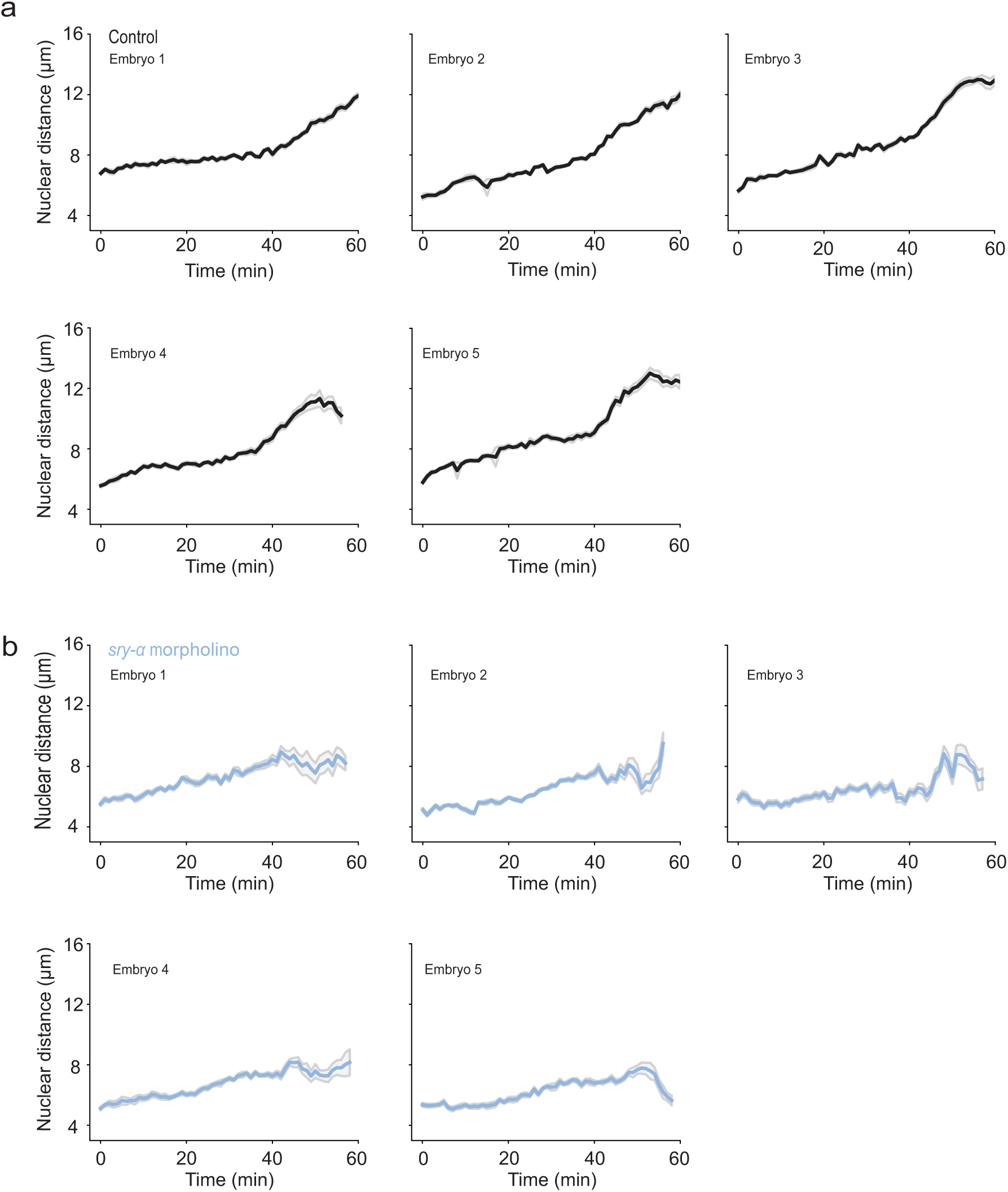
**a**,**b**, Individual embryo, raw nuclear distances for control morpholino (Control, black, **a**) and *sry-α* morpholino (*sry-α* morpholino, blue, **b**) injected embryos over the course of cellularization (15 nuclei followed per embryo; mean ± s.e.m. demarcated in gray).

**Extended Data Fig. 7 | Related to Fig. 6.**
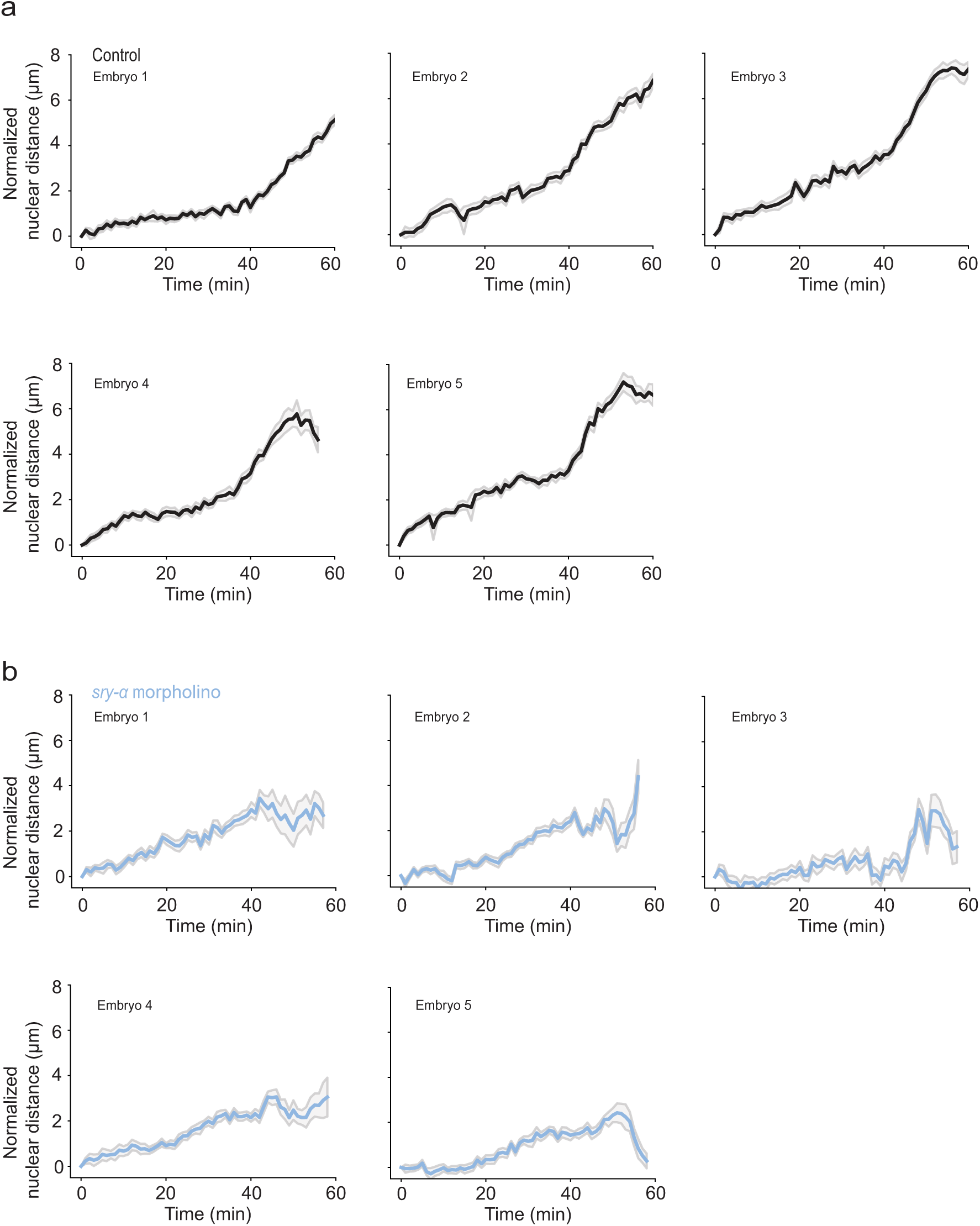
**a**,**b**, Individual embryo, normalized nuclear distances for control morpholino (Control, black, **a**) and *sry-α* morpholino (*sry-α* morpholino, blue, **b**) injected embryos over the course of cellularization (15 nuclei followed per embryo; mean ± s.e.m. demarcated in gray).

**Extended Data Video 1 (Related to Fig. 4):** Time lapse movie showing nuclear repositioning over the course of cellularization in a buffer-injected embryo (Control). Time is shown in minutes. 1 minute indicates the beginning of cellularization. Nuclei (magenta) elongate and move to a more basal position as cellularization proceeds. The vitelline membrane (orange line) serves as a marker for the embryo surface. Arrow shows relative movement of a single nucleus.

**Extended Data Video 2 (Related to Fig. 4):** Time lapse movie showing nuclear repositioning over the course of cellularization in a *sry-α* dsRNA-injected embryo (*sry-α* RNAi). Time is shown in minutes. 1 minute indicates the beginning of cellularization. Nuclei (magenta) elongate but fail to move to a more basal position as cellularization proceeds. Some nuclei move basally but reverse direction and “pop-up” to a very apical position. Some nuclei also tilt such that their long axis no longer remains at the normal 90° orientation to the cell surface. The vitelline membrane (orange line) serves as a marker for the embryo surface. Arrow shows relative movement of a single nucleus. Asterisks mark nuclei that “pop-up”.

**Extended Data Video 3 (Related to Fig. 6):** Time lapse movie showing nuclear repositioning over the course of cellularization in a control morpholino injected embryo (Control). Time is shown in minutes. 1 minute indicates the beginning of cellularization. Nuclei (magenta) elongate and move to a more basal position as cellularization proceeds. The vitelline membrane (orange line) serves as a marker for the embryo surface. Arrow shows relative movement of a single nucleus.

**Extended Data Video 4 (Related to Fig. 6):** Time lapse movie showing nuclear repositioning over the course of cellularization in a *sry-α* morpholino injected embryo (*sry-α* morpholino). Time is shown in minutes. 1 minute indicates the beginning of cellularization. Nuclei (magenta) elongate but fail to move to a more basal position as cellularization proceeds. Some nuclei move basally but reverse direction and “pop-up” to a very apical position. The vitelline membrane (orange line) serves as a marker for the embryo surface. Arrow shows relative movement of a single nucleus. Asterisks mark nuclei that “pop-up”.

