## Supplementary material for "Cytoplasmic localization of the mRNA encoding actin regulator, Serendipity-α, promotes adherens junction assembly and nuclear repositioning": All extended data referenced in the main manuscript

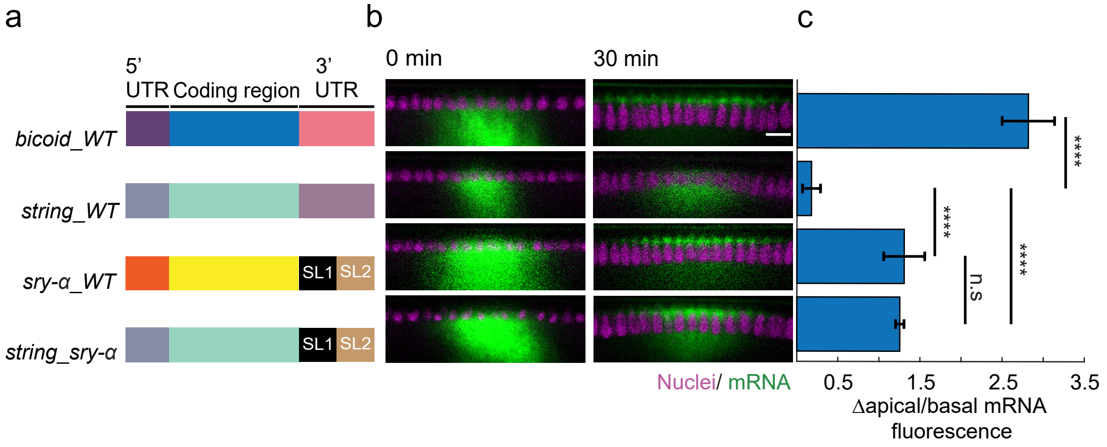
a
b
c
5' UTR
Coding region
3' UTR
bicoid_WT
string_WT
sry-α_WT
string_sry-α
SL1
SL2
0 min
30 min
Nuclei/ mRNA
****
****
****
n.s
0.5
1.5
2.5
3.5
Δapical/basal mRNA fluorescence

**Extended Data Fig. 1 | Related to Fig. 3e-g**

**a**, Schematic shows construct design for *in vitro* transcribed mRNAs for positive control (*bicoid_WT*), negative control (*string_WT*) and *sry-α* experimentals (*sry-α_WT*, *string_sry-α*; WT, wild-type; SL, stem loop). **b**, From time-lapse imaging, cross-sections show injected mRNA (green) and nuclei (Histone-mCherry, magenta). Identity of the mRNA is indicated by aligned construct in (**a**). Imaging started immediately after injection and proceeded for 30 minutes (0 min and 30 min, respectively). Scale bar = 10 μm. **c**, Change in apical to basal fluorescence from 0 to 30 minutes for the mRNA indicated by aligned construct in (**a**). Bars indicate mean ± s.e.m. (n = 5 embryos per construct); $p > 0.05$, not significant (n.s.); ****$p < 0.00005$, one-way ANOVA.

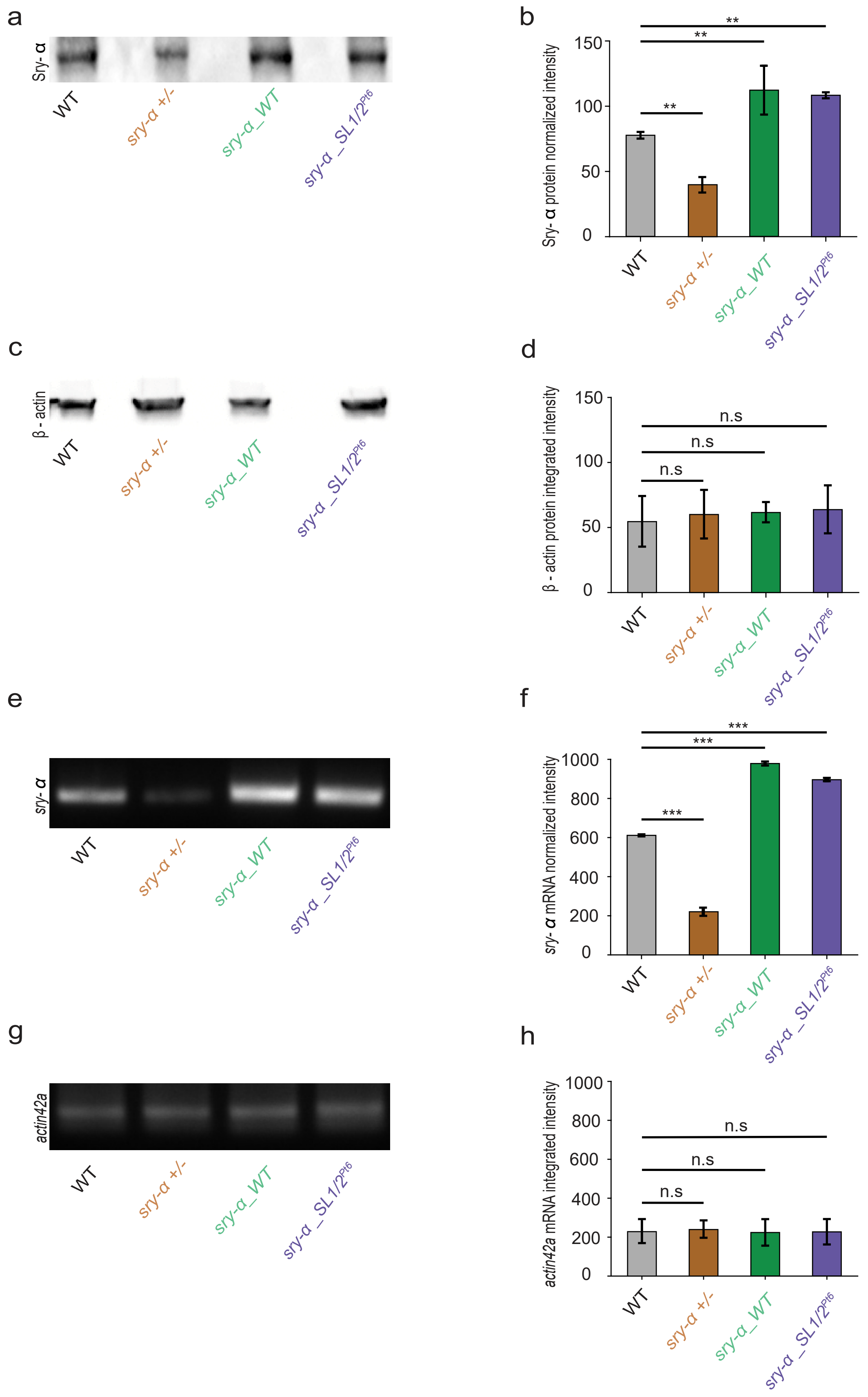
a
Sry- α
WT
sry-α +/-
sry-α _WT
sry-α _SL1/2Pt6
b
Sry- α protein normalized intensity
150
100
50
0
**
**
**
WT
sry-α +/-
sry-α _WT
sry-α _SL1/2Pt6
c
β - actin
WT
sry-α +/-
sry-α _WT
sry-α _SL1/2Pt6
d
β - actin protein integrated intensity
150
100
50
0
n.s
n.s
n.s
WT
sry-α +/-
sry-α _WT
sry-α _SL1/2Pt6
e
sry- α
WT
sry-α +/-
sry-α _WT
sry-α _SL1/2Pt6
f
sry- α mRNA normalized intensity
1000
800
600
400
200
0
***
***
***
WT
sry-α +/-
sry-α _WT
sry-α _SL1/2Pt6
g
actin42a
WT
sry-α +/-
sry-α _WT
sry-α _SL1/2Pt6
h
actin42a mRNA integrated intensity
1000
800
600
400
200
0
n.s
n.s
n.s
WT
sry-α +/-
sry-α _WT
sry-α _SL1/2Pt6

**Extended Data Fig. 2 | Related to Fig. 3h, i**

**a**,**c**, Immunoblots show Sry-α and β-actin bands, respectively, for protein lysates from embryos from indicated stocks. **b**,**d**, Quantification of Sry-α and β-actin protein levels, respectively, as determined by densitometry. **e**,**g**, Ethidium bromide stained agarose gels show reverse transcription-PCR products for *sry-α* and *actin42a*, respectively, following mRNA extraction from embryos from indicated stocks. **f**,**h**, Quantification of *sry-α* and *actin42a* PCR product levels, respectively, as determined by densitometry. PCR product levels are considered a proxy for mRNA levels in embryos.

**a**-**h**, Stocks are: OreR (wild-type, WT); *sry-α - /TM3Sb, hb::LacZ (sry-α +/-)*; *sry-α_WT; sry-α_SL1/2*$^{Pt6}$.

**b**,**d**,**f**,**h**, Bars indicate mean ± s.e.m. (n = 3 biological replicates); $p > 0.05$, not significant (n.s.); $^{**}p < 0.005$, $^{***}p < 0.0005$, $^{****}p < 0.00005$, Students t-test.

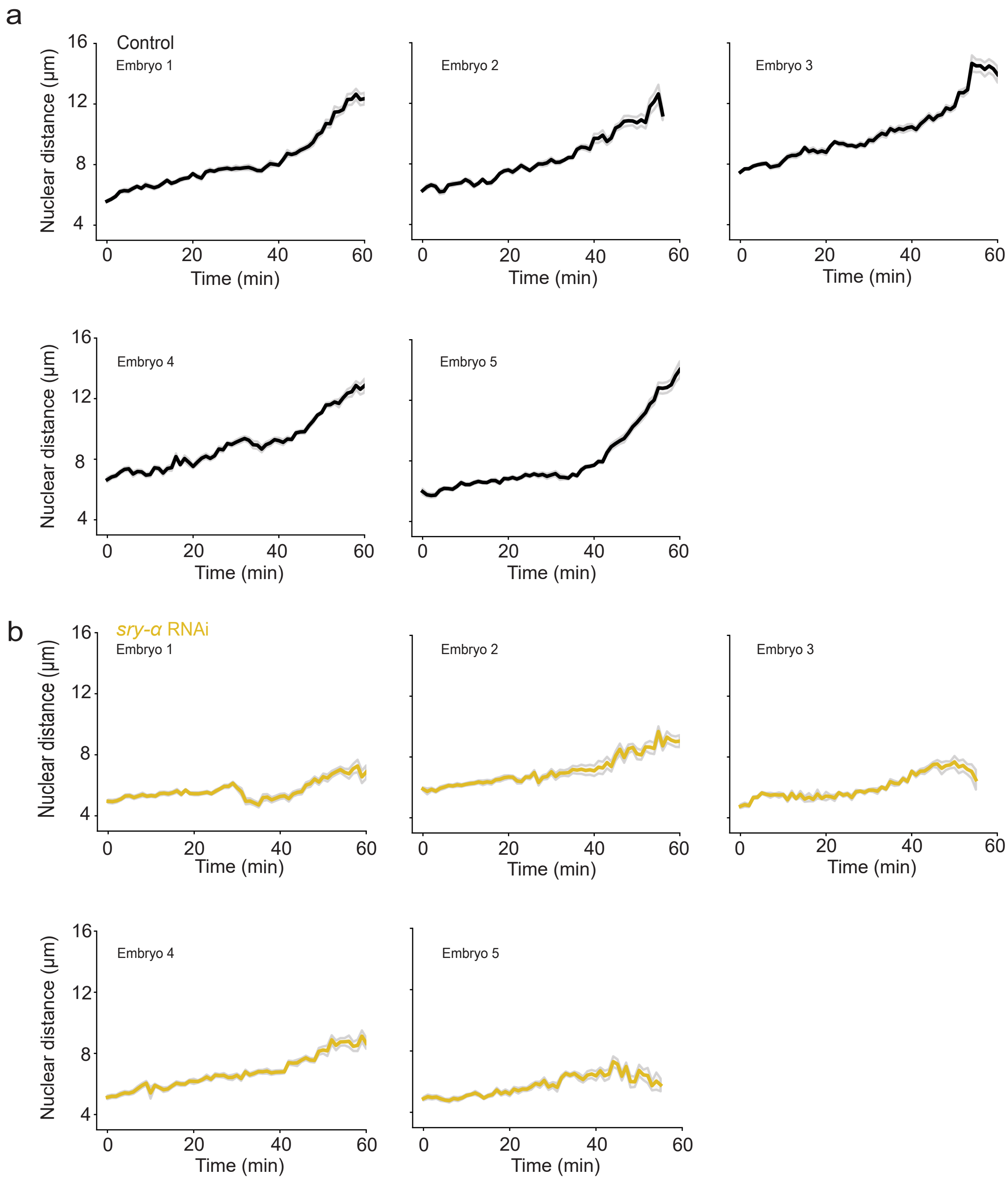

a
Control
Embryo 1
Embryo 2
Embryo 3
Embryo 4
Embryo 5
Nuclear distance (µm)
Time (min)
b
*sry-α* RNAi
Embryo 1
Embryo 2
Embryo 3
Embryo 4
Embryo 5
Nuclear distance (µm)
Time (min)

**Extended Data Fig. 3 | Related to Fig. 4**

**a**,**b**, Individual embryo, raw nuclear distances for buffer (Control, black, **a**) and *sry-α* dsRNA (*sry-α* RNAi, gold, **b**) injected embryos over the course of cellularization (15 nuclei followed per embryo; mean ± s.e.m. demarcated in gray).

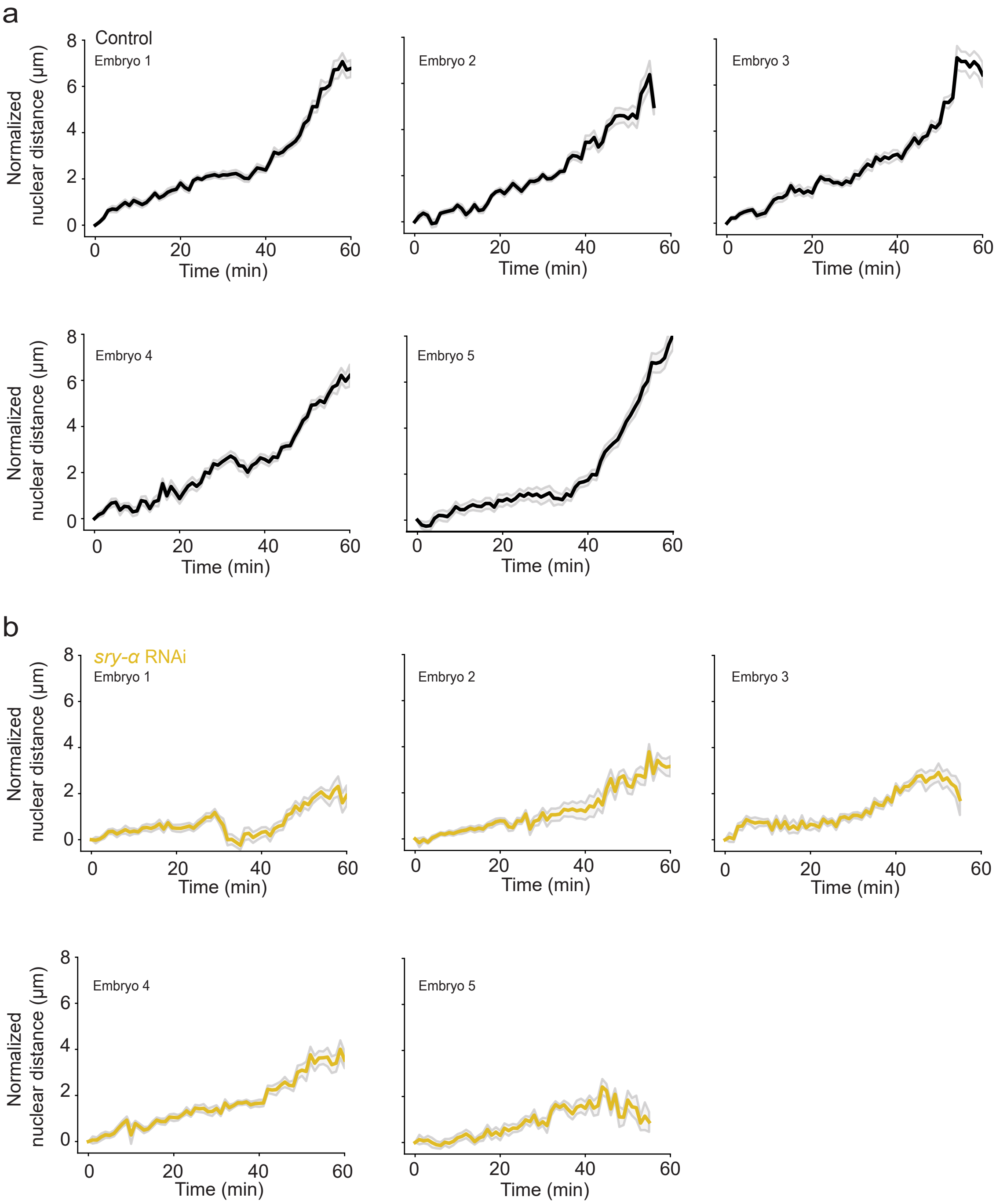

a
Control
Embryo 1
Embryo 2
Embryo 3
Embryo 4
Embryo 5
Normalized nuclear distance (µm)
Time (min)
0
2
4
6
8
0
20
40
60
b
sry-α RNAi
Embryo 1
Embryo 2
Embryo 3
Embryo 4
Embryo 5
Normalized nuclear distance (µm)
Time (min)

**Extended Data Fig. 4 | Related to Fig. 4**

**a**,**b**, Individual embryo, normalized nuclear distances for buffer (Control, black, **a**) and *sry-α* dsRNA (*sry-α* RNAi, gold, **b**) injected embryos over the course of cellularization (15 nuclei followed per embryo; mean ± s.e.m. demarcated in gray).

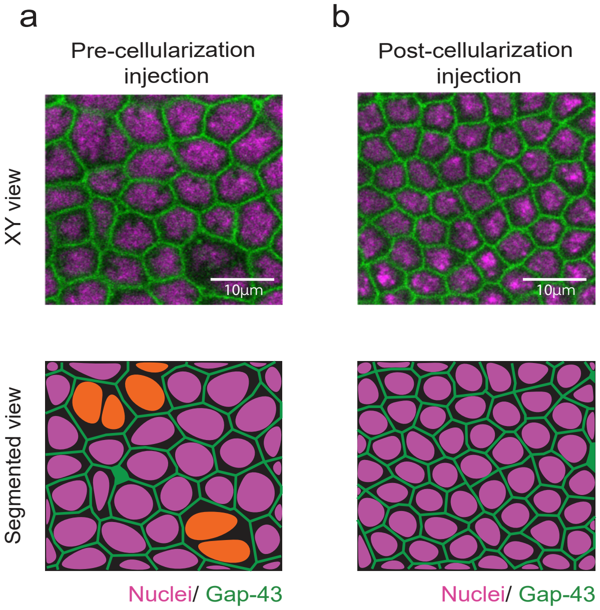

a
b
Pre-cellularization injection
Post-cellularization injection
XY view
Segmented view
10µm
10µm
Nuclei/ Gap-43
Nuclei/ Gap-43

**Extended Data Fig. 5 | Related to Fig. 6**

**a**,**b**, From time-lapse imaging, single plane confocal XY views show cellularization furrows (Gap43-GFP / plasma membrane, green) ingressing between nuclei (Histone-mCherry, magenta) in *sry-α* morpholino injected embryos. Embryos were either injected prior to cellularization (**a**) or just after cellularization onset (**b**; n = 5 embryos per treatment). Bottom row shows segmented views of the corresponding XY views with nuclei highlighted (orange, **a**) in multinucleated cells where furrows regressed. Images collected from embryos at late cellularization, with furrow lengths > 5 $\mu$m. Scale bars = 10 $\mu$m.

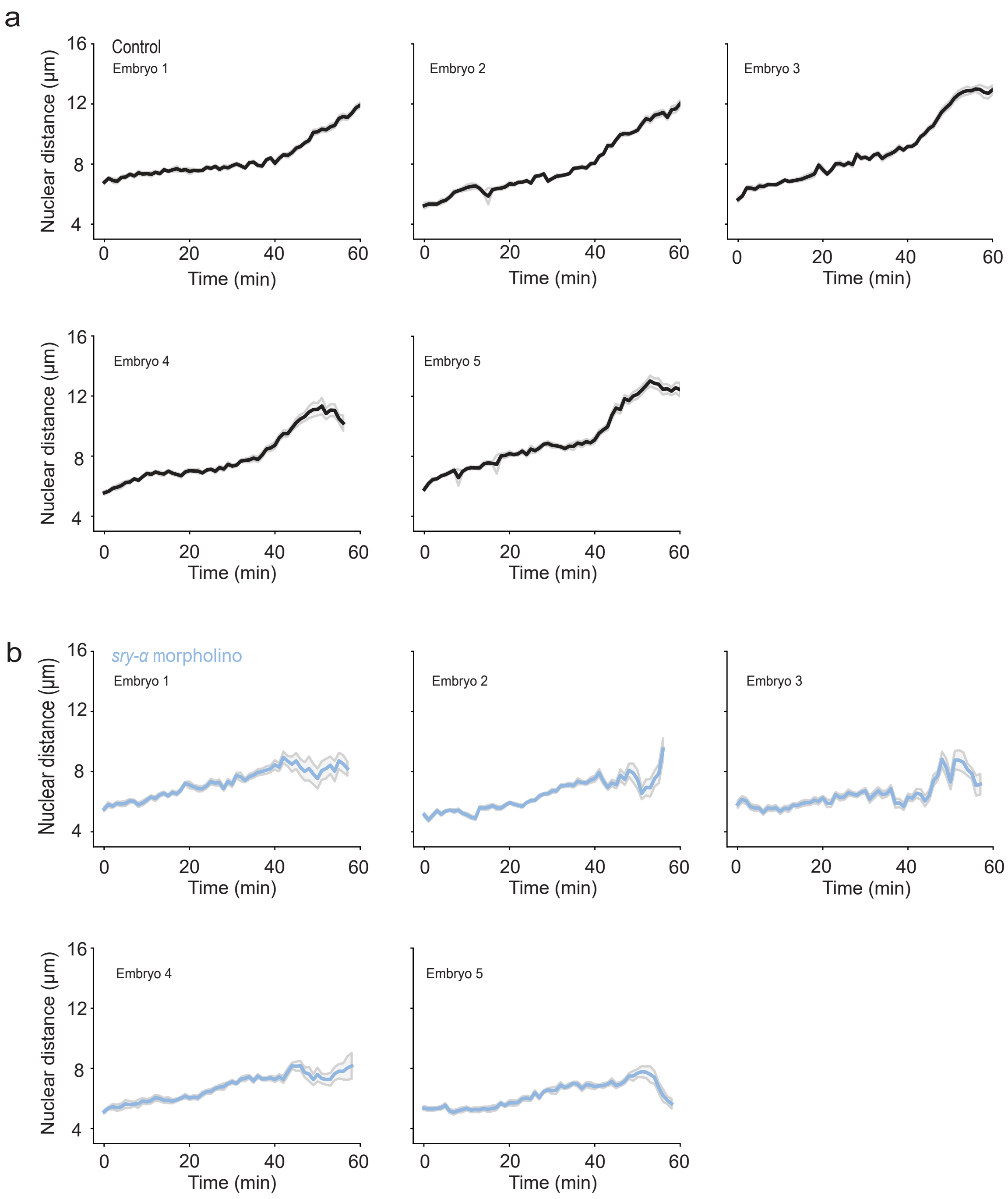
a
Control
Embryo 1
Embryo 2
Embryo 3
Embryo 4
Embryo 5
Nuclear distance (μm)
Time (min)
b
sry-α morpholino
Embryo 1
Embryo 2
Embryo 3
Embryo 4
Embryo 5
Nuclear distance (μm)
Time (min)

**Extended Data Fig. 6 | Related to Fig. 6**

**a**,**b**, Individual embryo, raw nuclear distances for control morpholino (Control, black, **a**) and *sry-α* morpholino (*sry-α* morpholino, blue, **b**) injected embryos over the course of cellularization (15 nuclei followed per embryo; mean $\pm$ s.e.m. demarcated in gray).

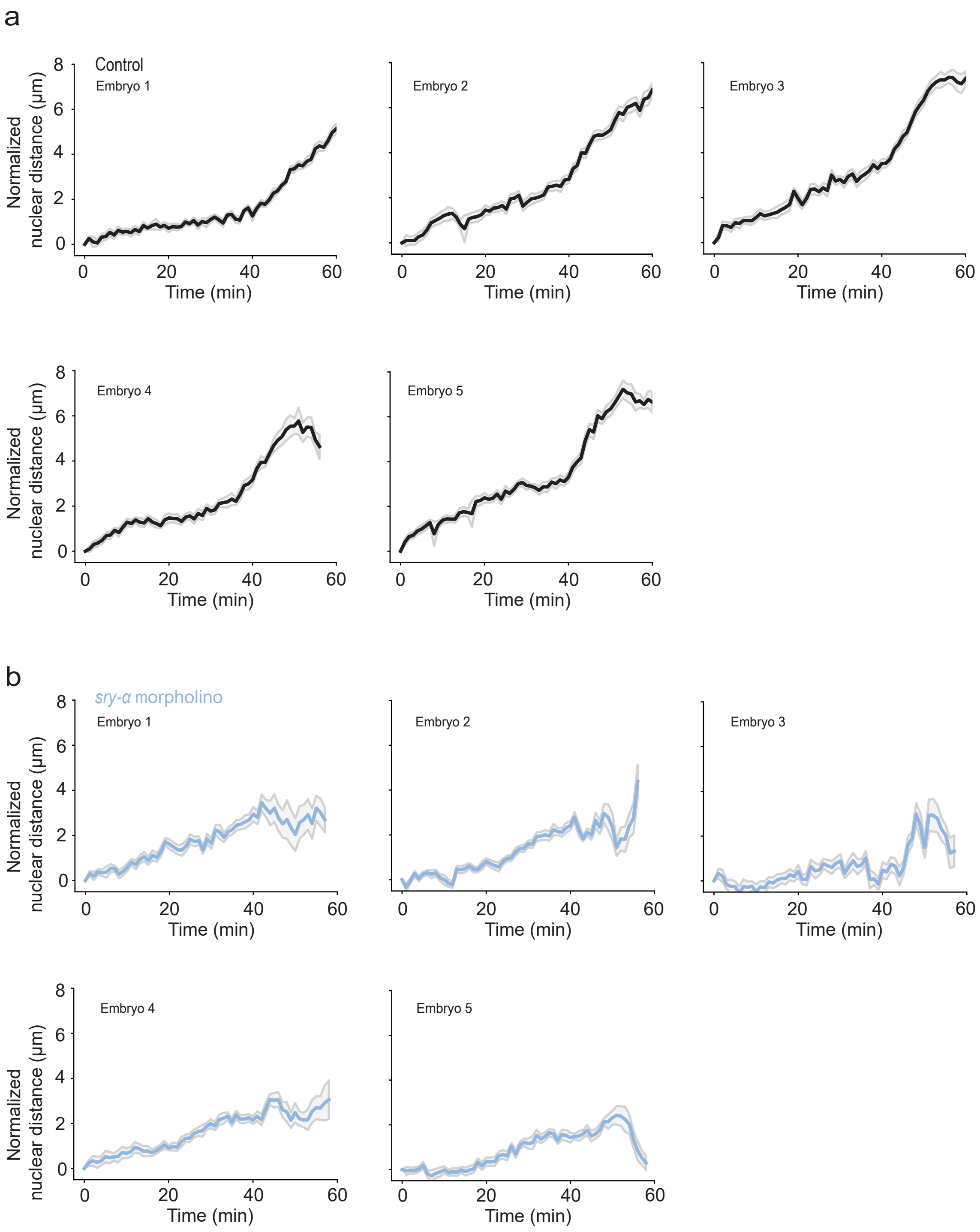
a
Control
Embryo 1
Embryo 2
Embryo 3
Embryo 4
Embryo 5
Normalized nuclear distance (µm)
Time (min)
b
*sry-α* morpholino
Embryo 1
Embryo 2
Embryo 3
Embryo 4
Embryo 5
Normalized nuclear distance (µm)
Time (min)

**Extended Data Fig. 7 | Related to Fig. 6**

**a**,**b**, Individual embryo, normalized nuclear distances for control morpholino (Control, black, **a**) and *sry-α* morpholino (*sry-α* morpholino, blue, **b**) injected embryos over the course of cellularization (15 nuclei followed per embryo; mean ± s.e.m. demarcated in gray).
